# GATA3 ZnF2-defective mutant condensation underlies type I IFN-activating in breast cancer

**DOI:** 10.1101/2023.05.02.538687

**Authors:** Yatao Chen, Yajie Wan, Xiaoying Pei, Tan Wang, Zhifang Ma, Liming Chen

## Abstract

Zinc finger (ZnF) transcription factors (TFs) consist of ZnF-containing DNA-binding domains (DBDs) and intrinsically disordered region (IDR)-containing activation domains (ADs). Recent studies have suggested that liquid-liquid phase separation (LLPS) is the fundamental mechanism underlying human health and disease, with ZnF TFs activating gene expression through the LLPS capacity of their IDR-containing ADs. However, little is known about how the well-folded DBD of ZnF TFs is involved in their LLPS mechanism. GATA3 is one of the most frequently mutated genes in breast cancer, and its encoded protein GATA3, which contains two ZnFs (ZnF1 and ZnF2) in its DBD, is a master regulator of immunity. Here, we show that GATA3 undergoes LLPS in cells and in vitro, and its DBD plays an important regulatory role. Mechanistically, ZnF2 in the DBD contains two arginine amino acids (R329 and R330) that provide critical charges to regulate GATA3 LLPS and DNA binding by generating multivalent electrostatic interactions. Functionally, we demonstrated that ZnF2-regulated GATA3 LLPS is the mechanism underlying the multifaceted function of GATA3 in breast cancer development and immune regulation, where aberrant GATA3 LLPS caused by artificial or breast cancer-associated ZnF2-defective mutations by reducing Suv39H1 protein stability showed significantly reduced potential in promoting breast cancer development and exhibited remarkably enhanced capacities for activating type I interferon signaling. Since ZnF is a common feature in the DBDs of ZnF TFs, by describing GATA3 as a proof-of-principle, our data suggest that ZnF-regulated LLPS may be a general mechanism underlying the multifaceted function of ZnF TFs in human health and disease.

## Introduction

Transcription factors (TFs) comprise of one or more well-folded DNA-binding domains (DBDs) and activation domains (ADs). The most prominent TFs in humans contain zinc fingers (ZnFs) in their DBDs, known as ZnF TFs, which play multifaceted functions in regulating various critical biological processes underlying human health and diseases (Cassandri et al., 2017; Jen and Wang, 2016). Recent evidence suggests that liquid-liquid phase separation (LLPS) is a phenotype-associated mechanism underlying human health and disease (Wang et al., 2021). The intrinsically disordered regions (IDRs) in protein sequences are considered the driving force for LLPS (Borcherds et al., 2021). Previous studies have shown that some Zn TFs, such as OCT4 and GCN4, activate their target genes through the phase separation capacity of their IDR-containing ADs (Boija et al., 2018). LLPS research is still in its infancy and further investigation is required to translate our current knowledge of LLPS into novel mechanistic, functional, and therapeutic discoveries. Little is known about how well-folded DBDs of ZnF TFs and ZnFs in their DBDs might be involved in their LLPS-mediated mechanisms underlying human health and disease, such as cancer.

The GATA TF family, an important Zn TF family consisting of six ZnF TFs (GATA1-6), has been well characterized to be involved in a variety of physiological and pathological processes, and many disease-associated mutations are located in and around the ZnF regions (Lentjes et al., 2016). GATA3, encoded by GATA3, is a typical member of the GATA TF family, and its DBD contains two ZnFs: ZnF1 and ZnF2. Studies in animal models and humans have characterized the important role of GATA3 in controlling the expression of a wide range of biologically and clinically important genes. Elevated GATA3 expression has been observed in various cancers, including breast cancer (Ikeda and Togashi, 2022). GATA3 is among the top three most frequently mutated genes in breast cancer (Ellis et al., 2012), and GATA3 is a master regulator of immunity (Wan, 2014). In this study, we investigated GATA3 to conceptually argue the role and mechanism of DBD and its ZnFs in the LLPS-mediated multifaceted function of ZnF TFs in human health and disease.

We found that GATA3 underwent LLPS both in vitro and in vitro. In GATA3 LLPS, its N-terminal domain (N), known as its AD, but not its C-terminal domain, is likely to play a driving role by containing IDRs, and its DBD, specifically ZnF2 but not ZnF1, plays an important regulatory role by containing two arginine amino acids: R329 and R330. In contrast to the IDRs in GATA3 AD, which contribute to LLPS through different types of multivalent interactions, ZnF2 in the GATA3 DBD regulates LLPS through multivalent electrostatic interactions (MEIs), with R329 and R330 in ZnF2 providing important charge blocks for binding with DNA. Aberrant LLPS caused by mutations in and around R329 and R330, including both artificial and breast cancer-associated mutations, showed reduced oncogenic potential in promoting breast tumor growth and enhanced the expression of immune genes in breast cancer cells, mainly interferon-stimulated genes (ISGs) for interferon (IFN) signaling. These data suggest that ZnF2 modulates the multifaceted function of GATA3 in breast cancer development and immune regulation by regulating LLPS of GATA3 by reducing Suv39H1 protein stability. Because charged amino acid-containing ZnFs with the intrinsic ability to generate MEIs are a common feature of ZnF TFs, GATA3 may serve as a proof-of-principle to suggest that LLPS regulated by ZnF-mediated MEIs is likely a general mechanism underlying the multifaceted function of ZnF TFs in many health and disease settings.

## Results

### GATA3 undergoes LLPS with chromatin in vivo

ZnF TFs on chromatin, which appear as vital and ubiquitous condensates in cell nuclei, are tightly and dynamically regulated to control specific gene expression and modulate many important phenotype-related biological processes (Voss and Hager, 2014). Indeed, an immunofluorescence (IF) assay revealed that GATA3 forms condensates on chromatin (DAPI indicates) without colocalization with non-chromatin condensation markers, including SC35, PML, and Colin (Fig. 1 A and B and fig. S1A). 1,6-hexanediol treatment and Fluorescence Recovery After Photobleaching (FRAP) assays have been widely used to assess whether protein condensates are driven by LLPS-mediated assemblies (Rippe, 2022). We found that GATA3-containing condensates on chromatin were disrupted upon 1,6-hexanediol treatment, and these condensates exhibited fluidity, as evidenced by fluorescence recovery after photobleaching in FRAP assays (Fig. 1 C-E). These results suggested the formation of GATA3-containing condensates on chromatin via the LLPS mechanism. GATA3-containing condensates on chromatin include many proteins besides GATA3. To test whether GATA3 undergoes LLPS for its condensation on chromatin, we performed in vitro LLPS assays using purified recombinant GATA3 protein, which is an MBP-GATA3 fusion protein that can be cleaved into GATA3 and MBP proteins by the PreScission HRV 3C protease. The results of the turbidity assay showed that the MBP-GATA3 protein cleaved by protease 3C exhibited remarkable and significantly increased turbidity compared to the control (Fig. 1F and fig. S1B). In an in vitro droplet formation assay, GATA3 with and without the mEGFP tag protein formed droplets (Fig. 1 G and H). Consistent with the observation in cells, 1,6-hexanediol treatment disrupted the droplets of purified GATA3-mGFP proteins, and the FRAP assay showed that the fluorescence of GATA3-mEGFP droplets can undergo fluorescence recovery after photobleaching (Fig. 1I and fig. S1C). Furthermore, a fusion event involving GATA3 droplets was observed (Fig. 1J). These results suggest that GATA3 has LLPS capacity for condensation on chromatin.

**Fig. 1.**
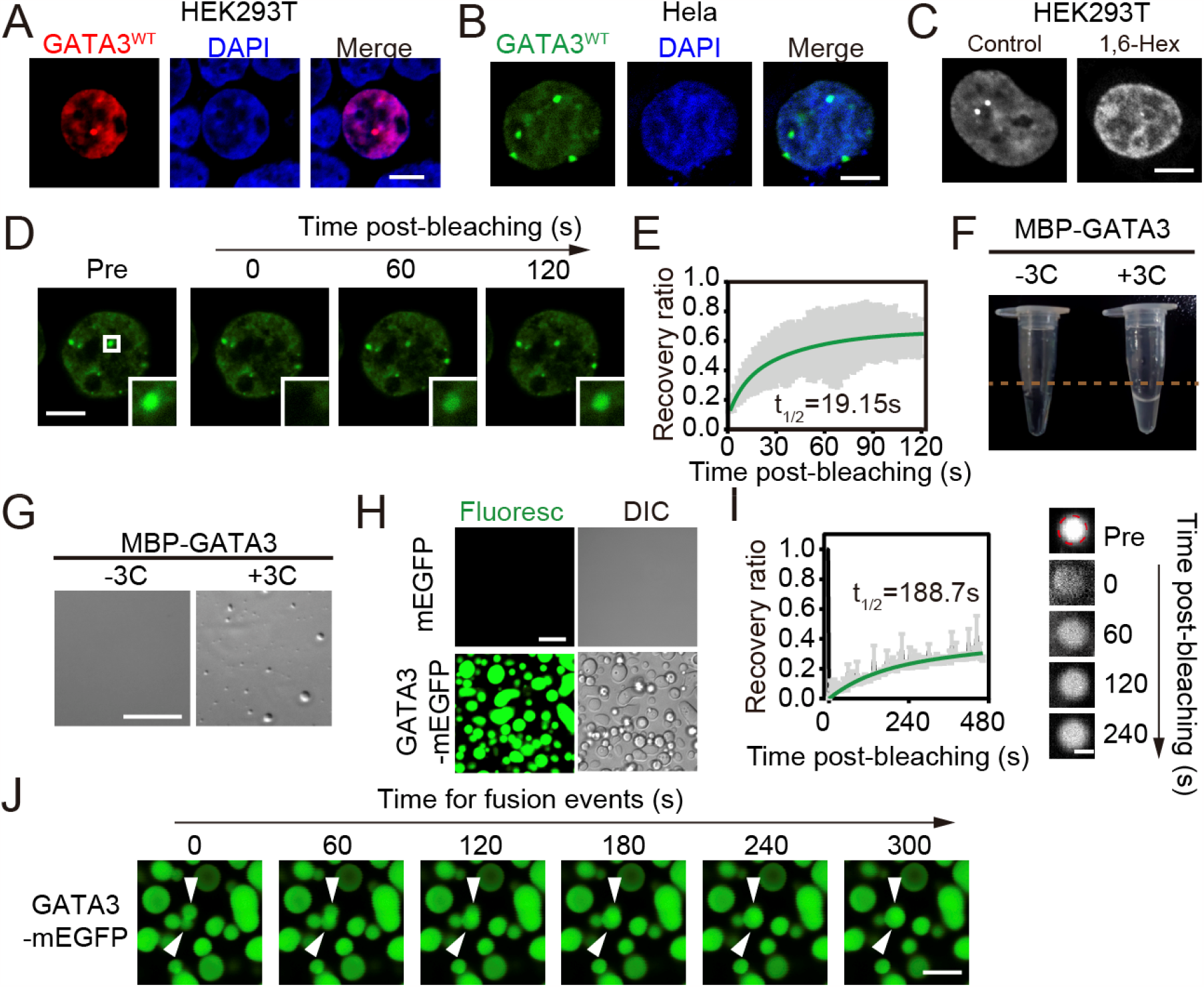
GATA3 undergoes LLPS in cells and in vitro. **(A)** Representative confocal microscopy images of HEK293T cells with ectopic expression of mCherry-tagged wild type GATA3 (GATA3^WT^). Scale bar, 5 μm. **(B)** Representative confocal microscopy images of Hela cells with ectopic expression of mEGFP-tagged GATA3^WT^. Scale bar, 2 μm. **(C)** Representative confocal microscopy images of HEK293T cells with ectopic expression of mEGFP-tagged GATA3^WT^ after treatment with 10% 1,6-hexanediol (1.6-Hex) compared to control. Scale bar,2 μm. **(D)** Representative images of mEGFP-tagged GATA3^WT^ droplets in HEK293T cells across time in FRAP assay. Scale bar, 2 μm. **(E)** Normalized fluorescence recovery of mEGFP-tagged GATA3^WT^ droplets in HEK293T cells as shown in (**D**), t_1/2_, means half-life of recovery time. **(F)** Representative images of 10 μM purified MBP-GATA3^WT^ protein without mEGFP tag in tubes treated with 3C protease treatment compared to control. **(G)** Representative images of droplets formed by 10 μM purified MBP-GATA3^WT^ protein without mEGFP tag with 3C protease treatment compared to control. Scale bar, 50 μm. **(H)** Representative images of droplets formed by 10 μM purified MBP-GATA3^WT^ protein with mEGFP tag compared to 10 μM purified MBP-mEGFP protein post the MBP cleaved by 3C protease treatment. Scale bar, 10 μm. **(I)** FRAP assay for droplets formed by 10 μM purified MBP-GATA3^WT^ protein with mEGFP tag after its MBP was cleaved by 3C protease treatment. Right, representative images of droplets across time in the FRAP assay. Light, normalized fluorescence recovery of droplets in FRAP assay. Scale bar, 1μm. **(J)** Representative images of droplet fusion events using 10 μM purified MBP-GATA3^WT^ protein with mEGFP tag after its MBP was cleaved by 3C protease treatment. Scale bar, 5 μm.

### LLPS of GATA3 is regulated by its structured DBD through multivalent electrostatic interactions

Zn TFs consist of structured DBDs containing ZnFs, and ADs containing IDRs. GATA3 has a DBD flanked by two IDR-containing domains: The N-terminal domain characterized as its AD and functionally unknown C-terminal domain (Fig. 2A). Some Zn TFs, such as OCT4 and GCN4, have been shown to activate their target genes through the phase-separating capacity of their IDR-containing Ads, but the role of DBD in LLPS behavior is largely unknown(Boija et al., 2018). Therefore, we investigated the role of the DBD domain in the LLPS of GATA3. In an in vitro droplet formation assay using purified protein, the number of droplets formed by a series of domain truncations, including the N-terminal domain (AD), DBD truncation (△DBD), C-terminal domain truncation (△C), and wild-type GATA3 (GATA3^WT^) (Fig. 2B). Our results suggest that the IDR-containing AD, but not C, is likely to provide the driving force, compared to GATA3^WT^, △DBD also showed reduced droplet formation ability (Fig. 2C), and only the DBD domain also has the ability of LLPS (fig. S2A and B). This indicates that DBD plays an enhancing role in GATA3 LLPS behavior in vitro.

**Fig. 2.**
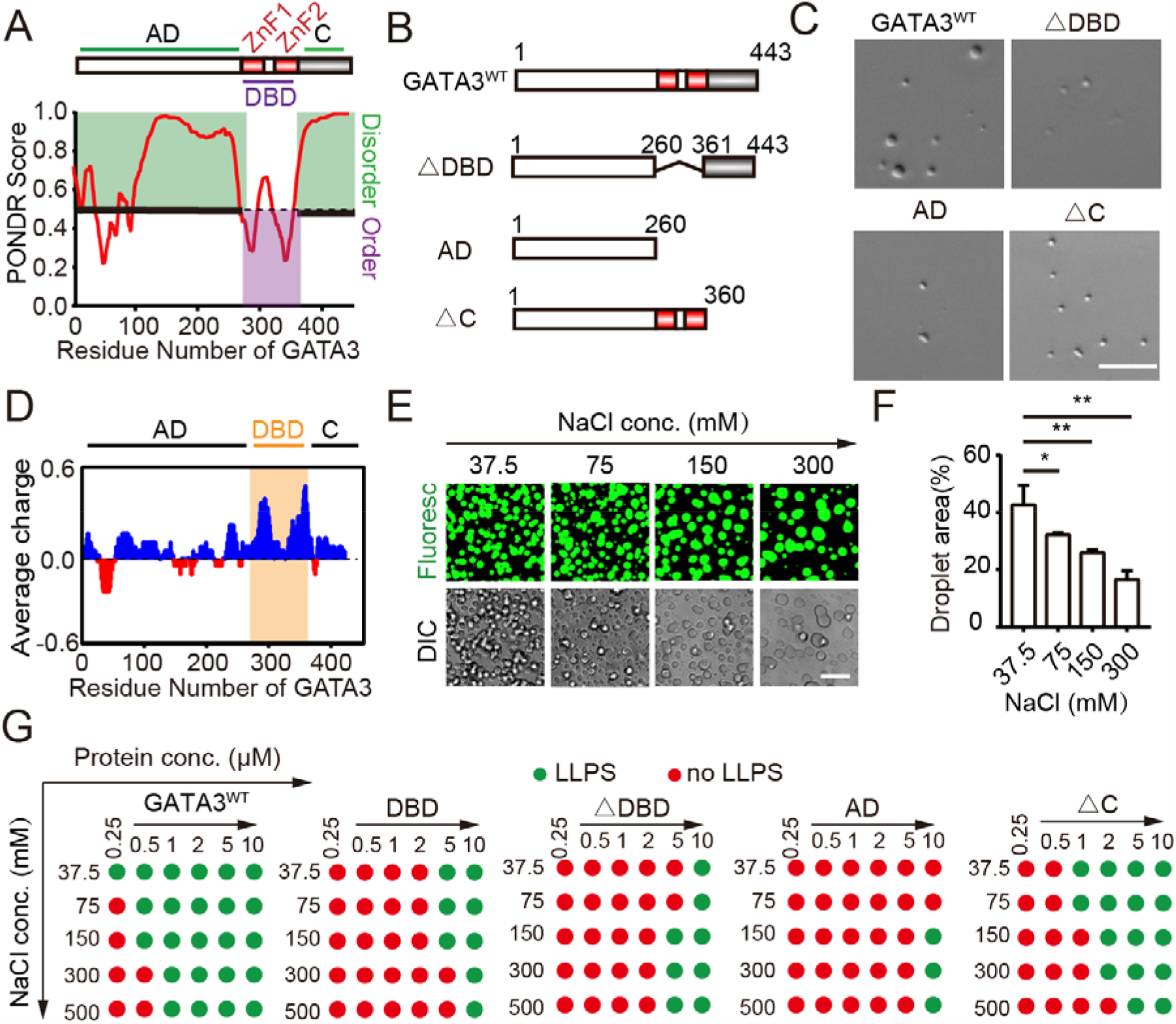
The DBD of GATA3 regulate phase separation via MEIs. **(A)** Disordered tendency analysis of GATA3 protein sequences. Top, a diagram shows the domain organization of GATA3. Bottom, PONDR assigned scores to the GATA3 protein sequence. A score higher than 0.5 indicates disordered regions (Green), and lower than 0.5 indicates ordered regions (Purple). **(B)** Diagrams for the domain structure of GATA3 truncations. **(C)** Representative images of droplets formed by 10 μM purified different MBP-GATA3 domain truncated proteins without mEGFP tag after cleavage of MBP by 3C protease treatment. Scale bar, 20 μm. **(D)** Average charge analysis of the GATA3 protein amino acid sequence. Red, negatively charged; blue, positively charged. Orange, DBD region. **(E)** Representative images of droplets formed by 5 μM GATA3-mEGFP with increasing NaCl concentration from 37.5 mM to 300 mM as indicated in the figure. Scale bar, 10 μm. **(F)** Quantification of the percentage of areas covered by droplets in the visual fields. Data are mean□±□s.d. *, p<0.05; **, p<0.01; ***, p<0.001(t-test). **(G)** Diagrams summarizing the LLPS of GATA3^WT^ and GATA3 domain truncated proteins with increasing of protein concentration from 0.25 μM to 10 μM along with increasing of NaCl concentration from 37.5mM to 500 mM as indicated in the figures. Green dots, LLPS; red dots, no LLPS.

Although not for Zn TFs, MEIs generated by charged amino acids have been characterized as another important source of force for the LLPS of several proteins(Rai et al., 2023). Charge analysis revealed a remarkably higher average charge in the DBD of GATA3 than in that of AD and C (Fig. 2D). MEI-mediated LLPS can be attenuated by increasing salt concentration(Yang et al., 2020). We found that GATA3^WT^ LLPS decreased with increasing NaCl concentrations (Fig. 2 E-F). In addition, the ability of GATA3^WT^, DBD, and △C (GATA3 mutants containing DBD) to form droplets decreased with increasing NaCl concentration. However, AD and △DBD did not (Fig. 2G). Furthermore, the fusion protein with ΔDBD-mEGFP and DBD-mEGFP also showed a difference in sensitivity with increasing NaCl concentration under the microscope (fig. S2C and D). Taken together, these results suggest that the structured DBD plays an important regulatory role in GATA3 LLPS via MEIs in vitro.

### ZnF2 R329 and R330 are essential to regulate GATA3 bind with DNA and LLPS behavior

We investigated the role of different domains in the LLPS of GATA3 in vivo. In both HEK293T and HeLa cells, we found that GATA3^WT^, AD, DBD, △DBD, △AD, and △C, but not C, formed granules in cells, with only AD and AD-containing △DBD forming significantly more granules in cell nuclei than GATA3^WT^ (Fig. 3 A and B and fig.S3A). This result also confirmed that AD and DBD play important roles in GATA3 LLPS in vivo. Are the differences in GATA3 truncation LLPS behavior in vitro and in vivo due to protein concentration? In cells, DBD truncations (including △DBD,C, and AD) showed higher protein expression compared to GATA3^WT^, △AD compared to C protein expression showed reduction, and △C protein expression showed reduction compared to AD (Fig. 3 C). This result suggests that the DBD may play an important regulatory role in GATA3 LLPS in vivo through protein concentration.

**Fig. 3.**
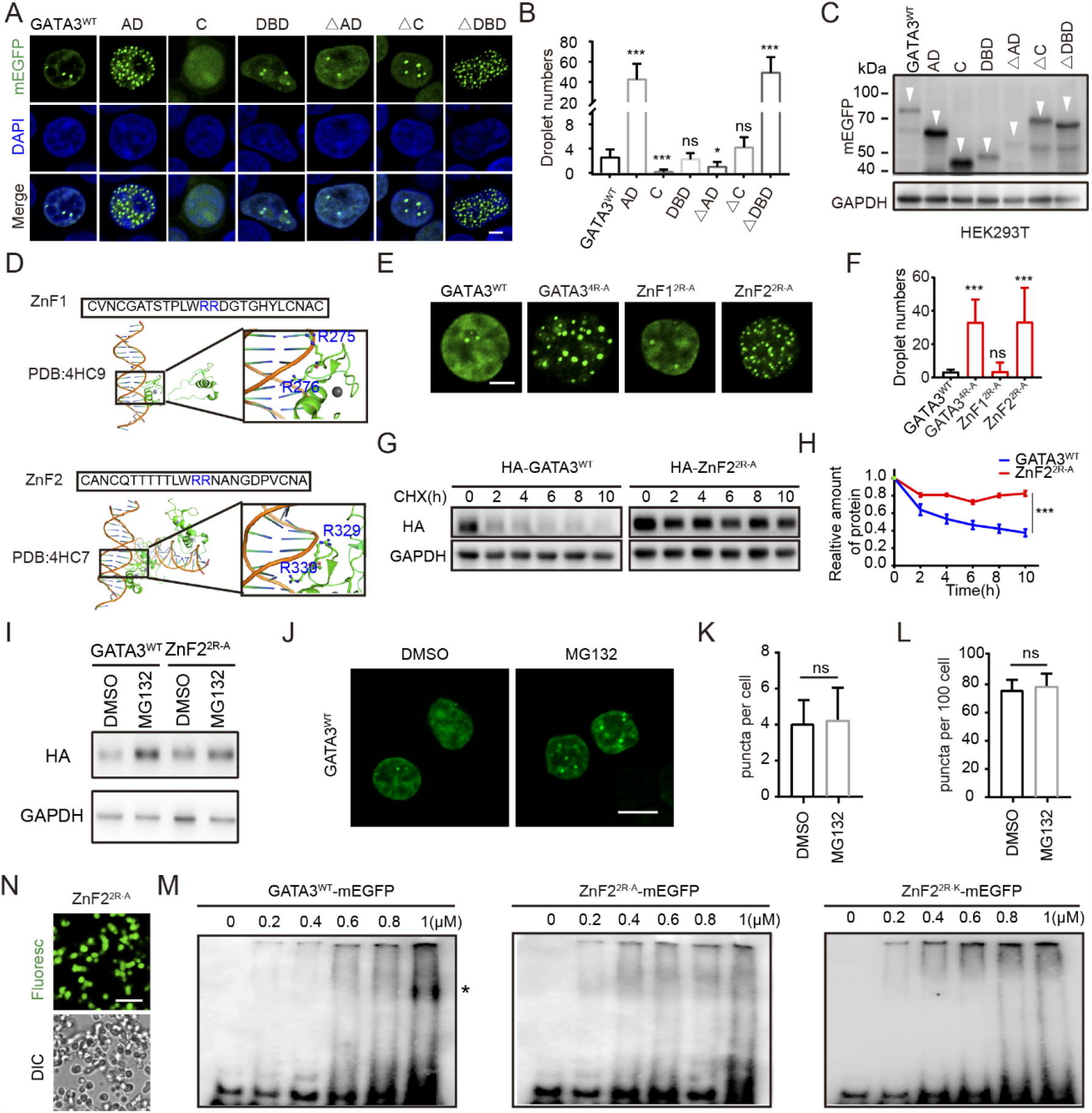
ZnF2 and its amino acids R329 and R330 regulate the LLPS of GATA3 and binding with DNA. **(A)** Representative confocal microscopy images of HEK293T cells with ectopic expression of mEGFP-tagged GATA3^WT^ and mEGFP-tagged truncations. Scale bar, 2 μm. **(B)** Quantification of the number of droplets per cell shown in (**A**). Biologically independent samples in triplicate: 20-50 transfected cells in each sample were quantified. Data are mean□±□s.d.*, p<0.05; **, p<0.01; ***, p<0.001 (t-test), ns, not significant. **(C)** Representative images of the mEGFP protein levels of mEGFP-tagged GATA3 truncations in HEK293T cells by Western blot. GAPDH was used as a loading control. White arrow indicates wild-type GATA3 and GATA3 truncations. **(D)** above, a representative image elucidating the complex structure of ZnF1 in GATA3 binding to DNA (PDB: 4HC9). below, a representative image elucidating the complex structure of ZnF2 in GATA3 binding to DNA (PDB: 4HC7). **(E)** Representative confocal microscopy images of HEK293T cells with ectopic expression of mEGFP-tagged GATA3 mutants compared to mEGFP-tagged GATA3^WT^ as indicated in the figure. Scale bar, 2 μm. **(F)** Quantification of the number of droplets per cell shown in (C). Data are mean□±□s.d. ***, p<0.001; ns, not significant (t-test). **(G)** Time analyzing of GATA3 protein expression in HEK293T cells transfected HA-GATA3^WT^ or HA-ZnF2^2R-A^ treated with CHX. **(H)** Statistical analysis of GATA3 protein expression amount treated with CHX (*p < 0.05; **p < 0.01; ***p < 0.001). **(I)** GATA3^WT^ or ZnF2^2R-A^ protein were analyzed in HEK293T cells treated with or without MG132(10 mM,8h). **(J)** Immunofluorescence images of HEK293T cells transfected GATA3^WT^, treated with or without MG132, scale bar, 5μm. **(K)** Quantification of the number of droplets per cell shown in (**J**). ns, not significant (t-test). **(L)** Quantification of the number of droplets positivity ratio cell shown in (**J**). ns, not significant (t-test). **(N)** Representative images of droplets formed by 10 μM purified MBP-ZnF2^2R-A^ protein with mEGFP tag after the MBP cleaved by 3C protease treatment. Scale bar, 10 μm. **(M)** Electrophoretic mobility shift assay of purified GATA3^WT^ and GATA3 mutants’ proteins showing biotinylated DNA as detected by horseradish peroxidase (HRP)-conjugated streptavidin. * indicates DNA-GATA3 complex.

Does DBD alter GATA3 LLPS behavior induced by MEI in vivo? GATA3 DBD with a known DNA-binding complex structure contains two ZnFs: ZnF1 and ZnF2 (Chen et al., 2012). ZnF1 and ZnF2 had the highest average charges in the DBD of GATA3, each containing two continuously charged arginine (R) amino acids (i.e., R^275^R^276^ in ZnF1 and R^329^R^330^ in ZnF2), which have been shown to bind to DNA in the complex structure (Fig.3 D). To investigate whether and how R^275^R^276^ in ZnF1 and R^329^R^330^ in ZnF2 affect GATA3 LLPS, we generated another set of GATA3 mutants: GATA3^4R-A^ with R^275^R^276^ in ZnF1 and R^329^R^330^ in ZnF2 mutated to alanine (A), ZnF1^2R-A^ with R^275^R^276^ in ZnF1 mutated to A^275^A^276^, and ZnF2^2R-A^ with R^329^R^330^ in ZnF2 mutated to A^329^A^330^. Interestingly, GATA3^4R-A^ and ZnF2^2R-A^ but not ZnF1^2R-A^ formed significantly more nuclear condensates in cells than GATA3^WT^ (Fig. 3 E and F). ZnF2^2R-A^ protein was more stable than GATA3^WT^ in cells (Fig. 3 G and H). Protein concentration is an important factor in the alteration of LLPS behavior; whether ZnF2^2R-A^ only influences GATA3 protein stability to affect GATA3 LLPS. When GATA3^WT^ protein expression increased after MG132 treatment, GATA3 condensation numbers and ratios in the nucleus did not change (Fig. 3 I -L). Consistently, ZnF2^2R-A^ showed altered LLPS behavior compared to GATA3^WT^ in an in vitro droplet formation assay (Fig. 1H, Fig. 3M, and fig. S3B). These results suggest that ZnF2^2R-A^ may affect the GATA3 ZnF2 structure by changing GATA3 LLPS behavior.

ZnF2, with its R329 and R330 residues, is essential for the regulation of DNA binding, and whether ZnF2^2R-A^ affects GATA3 binding with DNA. The EMSA results showed that, compared with GATA3WT, the ability of ZnF2^2R-A^ to bind with DNA was reduced. Furthermore, to verify the importance of ZnF2 with R329 and R330 MEI, we mutated R329 and R330 to lysine(K)-ZnF2^2R-K^, which provides a positive charge similar to R. The results showed that ZnF2^2R-K^, similar to ZnF2^2R-A^, displayed a decreased ability to bind with DNA and an increased ability to form condensates in the cell (Fig. 3N, and Fig. S3C). Taken together, these results suggest that ZnF2, but not ZnF1, is responsible for the regulatory role of GATA3 condensation, where R329 and R330 in ZnF2 are essential and irreplaceable amino acids that regulate DNA binding and GATA3 condensation.

### GATA3 binding with DNA regulates droplets’ dynamic properties in vivo and in vitro

ZnF2 of GATA3 is characterized by its specific binding to DNA containing GATA elements (Kumar et al., 2020). We have confirmed that ZnF2 with R329 and R330 positive charge forces is essential for binding with DNA and regulating GATA3 LLPS behavior, and whether ZnF2 binding with DNA also affects GATA3 droplet properties. In the in vitro droplet formation assay, we found that DNA containing GATA elements (GATA-DNA), but not DNA without GATA elements (noGATA-DNA), significantly altered the properties of GATA3 droplets. When GATA-DNA, which affects GATA3 MEI by binding to R329R330, GATA3 droplets showed reduced size and changed shape (Fig. 4 A and B). In comparison, both GATA-DNA and noGATA-DNA could alter the droplet size and shape of ZnF2^2R-A^, while the change was not significantly similar to GATA3(Fig. 4 C and D). Furthermore, FRAP assay results showed that GATA-DNA increased the fluidity of GATA3WT droplets, but not of ZnF2^2R-A^ droplets, compared to noGATA-DNA (Fig. 4 E and F). This result indicates that GATA3 relies on binding to GATA-DNA to control droplet formation and dynamic properties, whereas ZnF2^2R-A^ loses this dynamic adjustment ability due to reduced binding to GATA-DNA. Although fusion events of ZnF2^2R-A^ condensates were observed in cells, the fluorescence recovery of ZnF2^2R-A^ condensates on chromatin was significantly reduced compared to GATA3^WT^ in the FRAP assay (Fig. 4 G and H and fig.S4A). Taken together, ZnF2, with its R329 and R330 residues binding to DNA, regulates the dynamic properties of the droplets.

**Fig. 4.**
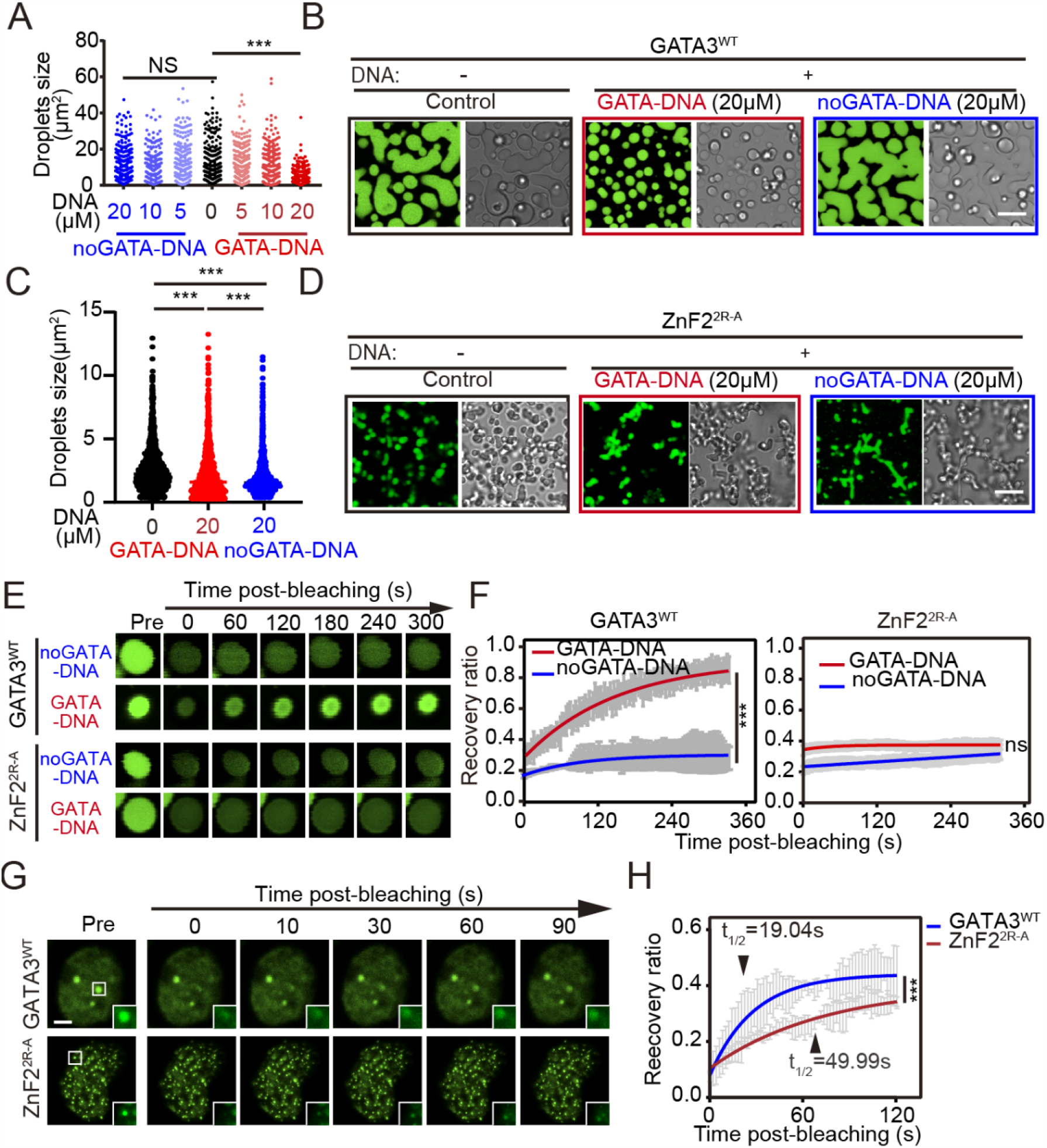
GATA3 binding with DNA regulates droplets’ dynamic properties in vivo and in vitro. **(A)** Quantification the size of droplets formed by 10 μM purified MBP-GATA3^WT^ protein with mEGFP tag and the MBP cleaved by 3C protease treatment in the visual fields, and DNA treatment was indicated in the figure. Blue, noGATA-DNA added. Red, GATA-DNA added. Black, no DNA added. GATA3-mEGFP protein concentration, 10 μM. **(B)** Representative images of droplets formed by 10 μM purified MBP-GATA3^WT^ protein with mEGFP tag and the MBP cleaved by 3C protease treatment with 20 μM noGATA-DNA or 20 μM GATA-DNA added compared to no DNA added. **(C)** Quantification the size of droplets formed by 10 μM purified MBP-ZnF2^2R-A^ protein with mEGFP tag and the MBP cleaved by 3C protease treatment in the visual fields, and DNA treatment was indicated in the figure. **(D)** Representative images of droplets formed by 10 μM purified MBP-ZnF2^2R-A^ protein with mEGFP tag and the MBP cleaved by 3C protease treatment with 20 μM noGATA-DNA or 20 μM GATA-DNA added compared to no DNA added. **(E)** Representative images of droplets formed by 10 μM purified mEGFP-tagged ZnF2^2R-A^ protein compared to 10 μM purified mEGFP-tagged GATA3^WT^ protein with 20 μM noGATA-DNA or 20 μM GATA-DNA added as indicated in the figure across time in FRAP assay. The MBP of purified proteins was cleaved by 3C protease treatment. Scale bars, 10 μm. **(F)** Normalized fluorescence recovery of droplets as shown in (**E**). Blue, 20 μM no-GATA DNA added. Red, 20 μM GATA DNA added. Black, no DNA added. **(G)** Representative images of mEGFP-tagged ZnF2^2R-A^ droplets compared to mEGFP-tagged GATA3^WT^ droplets in HEK293T cells across time in FRAP assay. Scale bars, 2 μm. **(H)** Normalized fluorescence recovery of mEGFP-tagged ZnF2^2R-A^(Red) droplets compared to mEGFP-tagged GATA3^WT^ (Blue) droplets in HEK293T cells as shown in (**G**), t_1/2_, half-life of recovery time.

### ZnF2-defective GATA3 mutants show reduced oncogenicity and aberrant LLPS

To explore the link between LLPS behavior and GATA3 function, we used breast cancer as an experimental model-GATA3 is one of the most frequently mutated genes in breast cancer (Ellis et al., 2012). Analysis of the gene mutation data archived in the Molecular Taxonomy of Breast Cancer International Consortium (METABRIC) revealed that in GATA3^mu^ proteins encoded by breast cancer-associated GATA3 mutations, ∼40% of GATA3^mu^ contain partially or completely defective ZnF2 (ZnF2^de^), ∼37% of GATA3^mu^ contain intact ZnF2 (ZnF2^in^), and the remaining ∼23% of GATA3^mu^ are highly variable (other mutations) (fig. S5A). To investigate whether breast-associated GATA3 mutants show aberrant LLPS behavior, four of the most frequently occurring mutations in breast cancer were selected for further investigation, including two ZnF2-defective GATA3 mutants (a splice mutation at X308-ZnF2^de-X308^ and a frameshift mutation at R330-ZnF2^de-R330^) and two ZnF2-intact GATA3 mutants (an M293K missense mutation-ZnF2^in-M293K^ and a G360 truncation mutation, ZnF2^in-G360^ (Fig. 5A). Similar to the artificial ZnF2-defective GATA3 mutant (ZnF2^2R-A^) and the artificial ZnF2-intact GATA3 mutant (ZnF1^2R-A^) described above, overexpression of breast cancer-associated ZnF2-defective GATA3 mutants (ZnF2^de-X308^ and ZnF2^de-R330^) but not breast cancer-associated ZnF2-intact GATA3 mutants (ZnF2^in-M293K^ and ZnF2^in-G360^) resulted in significantly more nuclear condensation compared to GATA3^WT^ in HEK293T cells (Fig. 5 B and C). ZnF2^de-R308^ condensation in cells could also be disassembled by 1,6-hexanediol treatment (Fig. 5D). Consistently, overexpression of ZnF2-defective GATA3 mutants (and ZnF2^de-R330^), but not ZnF2-intact GATA3 mutants (ZnF2^in-M293K^ and ZnF2^in-G360^), resulted in the formation of significantly more nuclear condensates compared to GATA3^WT^ in breast cancer cell lines, including T47D and MCF7^GATA3-KO^ (Fig. 5E and fig. S5 B and C). These results suggest that ZnF2-regulated GATA3 LLPS is involved in the development of breast cancer. To validate this hypothesis, we first examined the relationship between breast cancer-associated GATA3 mutants and the disease course of breast cancer. Consistent with previous observations, breast cancer patients with GATA3^mu^ showed a better overall prognosis in ERα-positive breast cancer patients than those with GATA3^WT^ (fig. S5D). Interestingly, breast cancer patients with ZnF2^de^ but not ZnF2^in^ mutations, showed a significantly better prognosis than those breast cancer patients with GATA3^WT^, suggesting that ZnF2^de^ is the major contributor to the better prognosis of patients with GATA3^mu^ versus GATA3^WT^ (Fig. 5F). We then examined the effects of ZnF2^de^ on the development of breast cancer and found that ZnF2-defective GATA3 mutants, including artificial ZnF2^2R-A^ and breast cancer-associated ZnF2^de-X308^ and ZnF2^de-R330^, showed reduced oncogenic capacity in promoting tumor growth in vitro and in vivo compared to GATA3^WT^ (Fig. 5 G-I). Although GATA3 is considered to play an important role in breast cancer development, its GATA3’s controversial role as an oncogene or tumor suppressor in breast cancer is still under debate. (Takaku et al., 2015) Both GATA3^mu^ and GATA3^WT^ are frequently upregulated in breast cancer (Asch-Kendrick and Cimino-Mathews, 2016). Our data suggest that elevated GATA3 in breast cancer, whether mutated or not, is oncogenic, while ZnF2-defective but not ZnF2-intact GATA3 mutants have significantly reduced oncogenic potential compared to wild-type GATA3 in promoting tumor growth by causing aberrant GATA3 LLPS, resulting in a better prognosis for patients with GATA3^mu^ compared to those with GATA3^WT^.

**Fig. 5.**
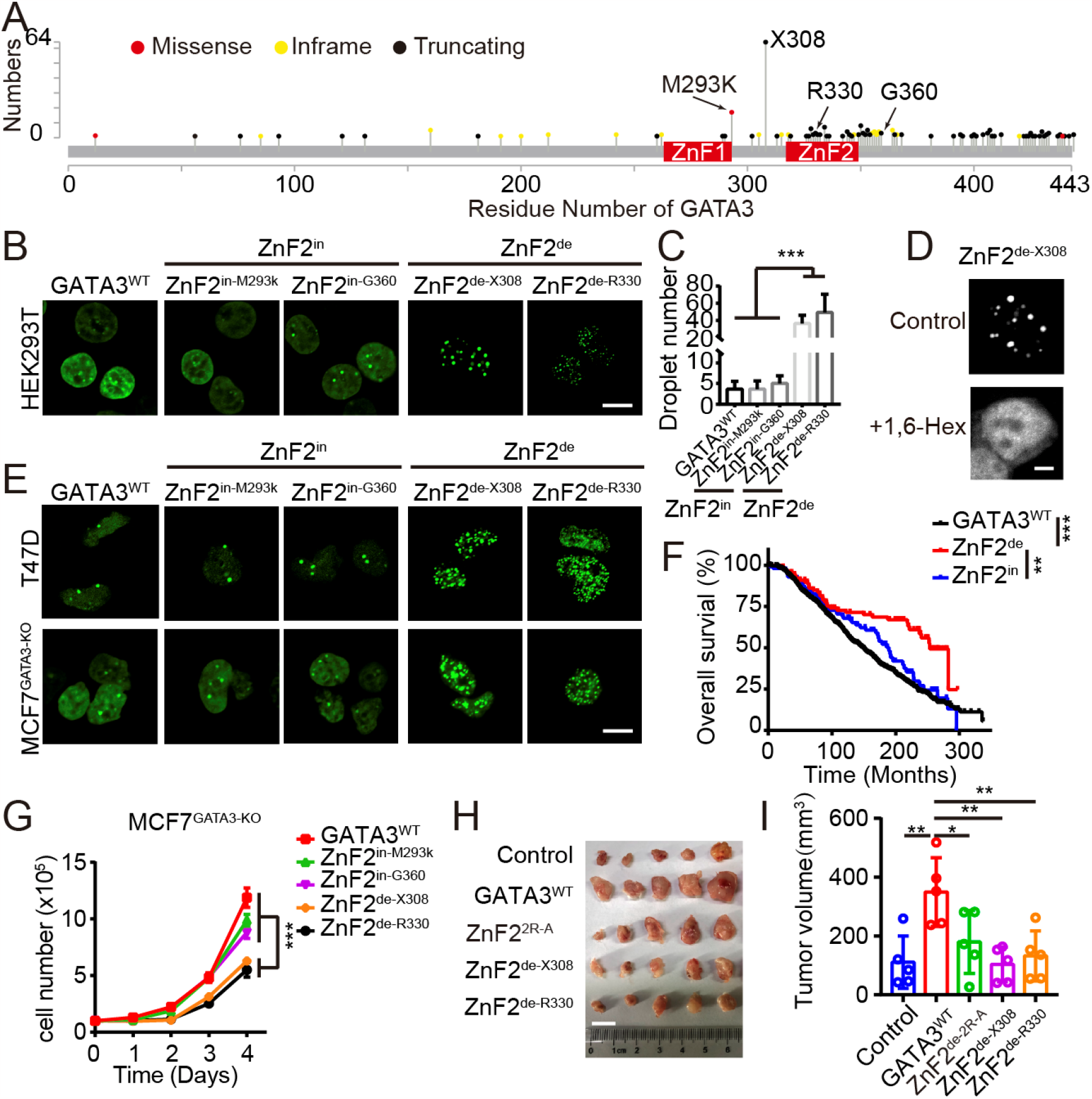
ZnF2-regulated LLPS of GATA3 unify explain the oncogenic role of GATA3 in breast cancer development. **(A)** A diagram summarizing the GATA3 mutations identified in clinical breast cancer samples. Black dots, GATA3 truncating mutations. Yellow dots, GATA3 in frame mutations. Red dots, GATA3 missense mutations. **(B**) Representative confocal microscopy images of HEK293T cells with ectopic expression of mEGFP-tagged breast cancer-associated GATA3 mutants compared to mEGFP-tagged GATA3^WT^ as indicated in the figure. Scale bar, 5□µm. **(C)** Quantification of the number of droplets per cell shown in (**B**). Data are mean ± s.d. ***, p<0.001 (t-test). **(D)** Images of HEK293T cells with ectopic expression of GATA3^de-X308^ genes and treatment with 10% 1,6-heleax indicated 1min. Scale bar, 2□µm. **(E)** Representative confocal microscopy images of breast cancer T47D and MCF7^GATA3-KO^ cells with ectopic expression of mEGFP-tagged breast cancer-associated GATA3 mutants compared to mEGFP-tagged GATA3^WT^ as indicated in the figure. MCF7^GATA3-KO^ is MCF7 with both endogenous GATA3^WT^ and GATA3^mu^ knocked out by Cas9. Scale bar, 5□µm. **(F)** Overall survival analysis of breast cancer patients with different types of mutations compared to those with wild type GATA3. GATA3^WT^(n=903): wild-type GATA3, ZnF2^de^(n=92): GATA3 mutations with defective ZnF2, ZnF2^in^(n=86): GATA3 mutations with intact ZnF2. **, p<0.01; ***, p<0.001 (log-rank p). **(G)** Growth potential of MCF7^GATA3-KO^ cells with ectopic overexpression expression of indicated GATA3 wild type or mutated genes in vitro by proliferation assay. Data are mean ± s.d. ***, p < 0.001 (t-test). **(H)** Growth potential of 4T1 cells with ectopic overexpression expression of indicated GATA3 wild type or mutated genes in vivo using xenograft breast cancer mouse model. Scale bar, 1□cm. **(I)** Quantification of tumor volumes in (**H**). Data are mean ± s.d. *, p < 0.05; **, p < 0.01 (t-test).

### ZnF2-defective GATA3 mutants enhances activation of IFN signaling

GATA3 has been characterized as a master immune regulator, as shown in many studies performed on immune cells (Wan, 2014). In addition to immune cells, an increasing number of studies have reported the essential roles of immune modulation in non-immune cells such as cancer cells (Dainichi et al., 2021). However, the function of GATA3 in non-immune cells remains largely unknown. To further argue that ZnF2-regulated GATA3 LLPS is an important mechanism underlying the multifaceted function of GATA3, we investigated whether and how aberrant ZnF2-regulated GATA3 LLPS is also relevant to immune events in breast cancer cells. To address this, we performed RNA-seq to examine the alteration of the transcriptional program by stable expression of ZnF2-defective GATA3 mutants, including artificial ZnF2^2R-A^ and disease-associated ZnF2^de-X308^ and ZnF2^de-R330^, compared to overexpression of GATA3^WT^, in MCF7^GATA3-KO^ breast cancer cells. Gene Ontology(GO) analysis of the genes differentially expressed in GATA3^WT^ cells compared with ZnF2^2R-A^ revealed a strong upregulation of apoptosis and immune-related hallmark gene sets (Fig. 6A). Gene set enrichment analysis (GSEA) of the genes differentially expressed in GATA3WT cells compared to ZnF2-defective GATA3 mutants revealed strong upregulation of type I IFN-related hallmark gene sets (Fig. 6B). Volcanic maps showed changes in these hallmark genes in ZnF2-defective GATA3 mutants compared with GATA3^WT^ cells (Fig. 6C). Among the differentially expressed genes between a ZnF2-defective GATA3 mutant and GATA3^WT^, 25 genes consistently showed differential expression in ZnF2^2R-A^, ZnF2^de-X308^ and ZnF2^de-R330^ compared to GATA3^WT^ (fig. s6A). Interestingly, 13 of these 25 genes were immune genes, all of which were significantly upregulated in the ZnF2-defective GATA3 mutants compared to GATA3^WT^ (Fig. 6D). Notably, 11 of the 13 immune-related genes were ISGs, including ISG15, IFI6, IFI27, IFI44L, IFIT1, IFIT3, IFI44, IFITM1, OAS2, OAS1, and OASL.

**Fig. 6.**
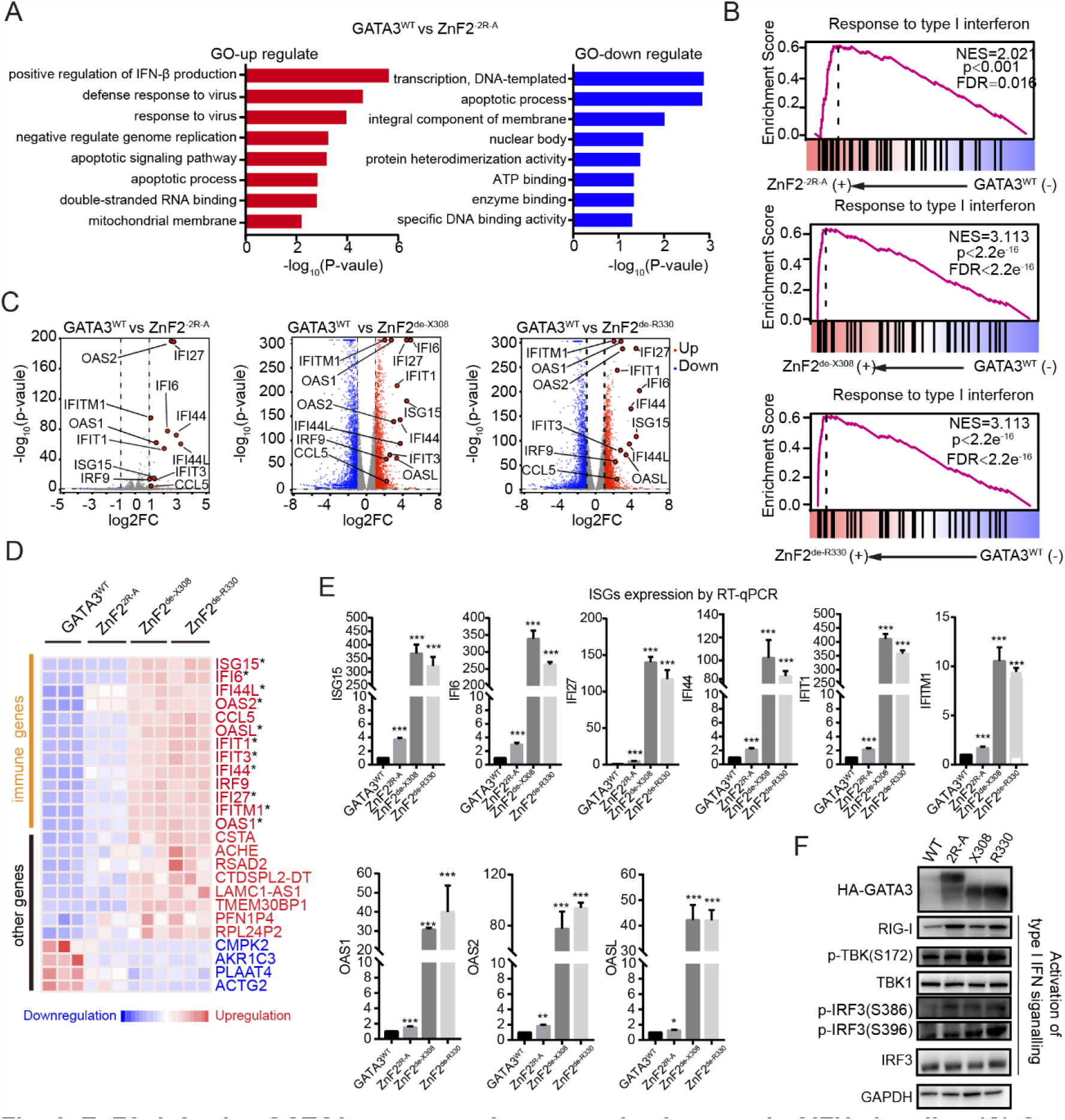
ZnF2-defective GATA3 mutants enhances activation type I of IFN signaling. **(A)** Gene ontology analysis for the indicated gene clusters from GATA3^WT^ VS ZnF2^2R-A^. **(B)** Diagrams show the impact ZnF2^2R-A^, ZnF2^de-X308^, and ZnF2 ^de-R330^ of versus GATA3^WT^ on the enrichment and upregulation of type I of IFN signaling in tumors via analyzing the RNA-seq data using GSEA. **(C)** Volcano plots representing the expression changes of all genes: significantly down-and up-regulated genes (GATA3^WT^ VS ZnF2^2R-A^, GATA3^WT^ VS ZnF2 ^de-X308^, GATA3^WT^ VS ZnF2 ^de-R330^: |log2 FC| >1 and p < 0.05) are colored blue and red, respectively; genes that do not show significant expression changes are colored gray; selected differentially expressed immune genes are labeled with gene names. **(D)** A heatmap depicting the common differently altered genes in ZnF2 defective mutants including ZnF2^2R-A^, ZnF2^de-X308^, and ZnF2 ^de-R330^ compared to GATA3^WT^, where upregulated genes are colored red, downregulated genes are colored blue, and “*” marks ISGs. **(E)** ISG15, IFI6, IFI27, IFI44, IFIT1, IFITM1, OAS1, OAS2 and OASL expression in MCF7^GATA3-KO^ with stable expression of ZnF2^2R-A^, ZnF2 ^de-X308^, and ZnF2 ^de-R330^ compared to GATA3^WT^ by RT-qPCR. Data are mean ± s.d. **, p < 0.01; ***, p < 0.001 (t-test). **(F)** Representative images of western blot analysis showing the level of IRF3, p-IRF3-S389/S396, TBK1 and p-TBK1-S172 in MCF7^GATA3-KO^ with stable expression of ZnF2^2R-A^, ZnF2^de-X308^, and ZnF2 ^de-R330^ compared to GATA3^WT^. GAPDH was used as an internal loading control.

Consistent with RNA-seq, RT-qPCR confirmed that all 11 ISGs were significantly upregulated by ZnF2-defective GATA3 mutants compared to GATA3^WT^, suggesting an important role of GATA3 in IFN signaling (Fig. 6E and fig. S6B). IRF3 and TBK1 are key regulators of type I IFN signaling, and their activation can be examined by the increased levels of IRF3-S386 and IRF3-S396 phosphorylation (p-IRF3-S386 and p-IRF3-S396) and TBK1 (p-TBK1) (Liu et al., 2015). We found that the stable expression of ZnF2-defective GATA3 mutants, such as ZnF2^de-X308^ and ZnF2^de-R330^, increased the levels of p-IRF3-S386, p-IRF3-S396, and p-TBK1 (Fig. 6F). In support of the more potent role of ZnF2-defective GATA3 mutants in activating type I IFN signaling compared to GATA3^WT^, the expression levels of two type I IFN genes, IFNA1 and IFNB1, were significantly increased in breast cancer cells expressing ZnF2^de-X308^ and ZnF2^de-R330^ compared to GATA3^WT^ cells, while ZnF2^2R-A^ not change compared to GATA3^WT^ cells (fig. S6C). These results suggest that ZnF2 and its R^329^R^330^ of GATA3 play an important role in innate immunity, specifically in the activation of IFN signaling, in which GATA3 LLPS behavior participates.

### GATA3 LLPS participates in Suv39H1-regulated chromatin subcompartments modulating type I IFN signal activation

It has been well characterized that ZnFs in the DBDs of ZnF TFs are essential for their binding to specific DNA elements in chromatin to localize to chromatin subcompartments (CSCs) to control specific gene expression. CSCs that are positive for transcriptional activation markers, such as p300, which is typically recruited to transcriptional enhancers, and H3K27ac, which is an enhancer marker, are often under active transcription (Loven et al., 2013; Ortega et al., 2018). In support of ZnF2-defective GATA3 mutants increasing the expression of ISGs compared to GATA3^WT^ described above, IF assays showed that in MCF7^GATA3-KO^ breast cancer cells, overexpression of ZnF2-defective GATA3 mutants, including artificial ZnF2^2R-A^ and breast cancer-associated ZnF2^de-X308^, increased their colocalization with p300-positive and H3K27ac-positive CSCs compared to GATA3^WT^ (Fig. 7 A). In contrast to p300 and H3K27ac, H3K9me3 is associated with silencing gene expression (Richards and Elgin, 2002). Consistently, IF assays showed that in MCF7^GATA3-KO^ breast cancer cells, the condensation of ZnF2-defective GATA3 mutants, including artificial ZnF2^2R-A^ and breast cancer-associated ZnF2^de-X308^, resulted in reduced H3K9me3 levels compared to GATA3^WT^ (Fig. 7 B). Previous studies have identified ∼3, 000 GATA3-bound genes with >14,000 GATA3 binding sites in the immune cells (Kanhere et al., 2012a; Voss and Hager, 2014; Wan, 2014). These GATA3-bound genes identified in immune cells include ISGs, such as IFIT3, OASL, IFIT1, IFI44L, IFI27, and OAS1, which are regulated by GATA3 in a ZnF2-mediated manner in breast cancer cells (Kanhere et al., 2012b). When we examined GATA3-bound genes in breast cancer cells with or without ZnF2-defective mutants by analyzing publicly available datasets, we found that in breast cancer cells, breast cancer-associated ZnF2-defective mutants, including ZnF2^de-X308^ and ZnF2^de-R330^, were remarkably more enriched in ISGs, such as IFI6 and IFI27(recognized by ZnF1 through GATC/G), compared to GATA3^WT^ (Fig. 7 C and fig. S7A). These results indicate that ZnF2-defective mutants bind to ISG genes to regulate gene expression. In addition, how does GATA3 access these genomic locations through LLPS? Suv39H1 and SETDB1 are major histone lysine methyltransferases that generate H3K9me3, which regulates the open/closed state of the genome (Padeken et al., 2022; Sanulli et al., 2019). In MCF7^GATA3-KO^ breast cancer cells, the expression of ZnF2-defective GATA3 mutants, including artificial ZnF2^2R-A^ and breast cancer-associated ZnF2^de-X308^ and ZnF2^de-R330^, decreased Suv39H1 but not SETDB1 protein levels, and reduced H3K9me3 levels compared to GATA3^WT^ (Fig. 7 D). Furthermore, ZnF2-defective GATA3 mutants did not significantly alter the mRNA expression level of Suv39H1 but significantly decreased Suv39H1 protein concentration compared to GATA3^WT^, suggesting that ZnF2-defective GATA3 mutants decreased Suv39H1 protein levels by destabilizing the Suv39H1 protein (Fig. 7 E-G). CSCs are formed via an LLPS-mediated mechanism, and Suv39H1 is important for HP1-H3K9me3-related CSCs formation(Rippe, 2022). Does Suv39H1 regulate CSC to permit GATA3 condensation formation In MCF7^GATA3-KO^ breast cancer cells expressing GATA3^WT^, treatment with Chaetocin, an inhibitor of Suv39H1 to disrupt H3K9me3 modification, increased GATA3 condensed numbers, and the expression of ZnF2 defective mutants regulated the above immune genes, which contains GATA3 bound ISGs, including IFIT3, OASL, IFIT1, IFI44L, IFI27, and OAS1, compared to the control (Fig. 7 H). Furthermore, ZnF2^2R-A^ and GATA3 colocalized with Suv39h1. However, compared to GATA3^WT^, ZnF2^2R-A^ showed reduced colocalization with HP1β because of Suv39h1 reduction. (fig. 7 B and C). These results suggest that Suv39H1-regulated CSC formation also contributes to GATA3 condensation, which affects ISG gene expression, even though the GATA3 ZnF2 structure is not mutated. It will be interesting to further investigate how ZnF2-defective GATA3 mutants destabilize Suv39H1 proteins to regulate LLPS-mediated formation of CSCs. Taken together, our results suggest that GATA3 ZnF2-regulated LLPS is an important mechanism underlying Suv39H1 in controlling immune gene expression for immune regulation.

**Fig. 7.**
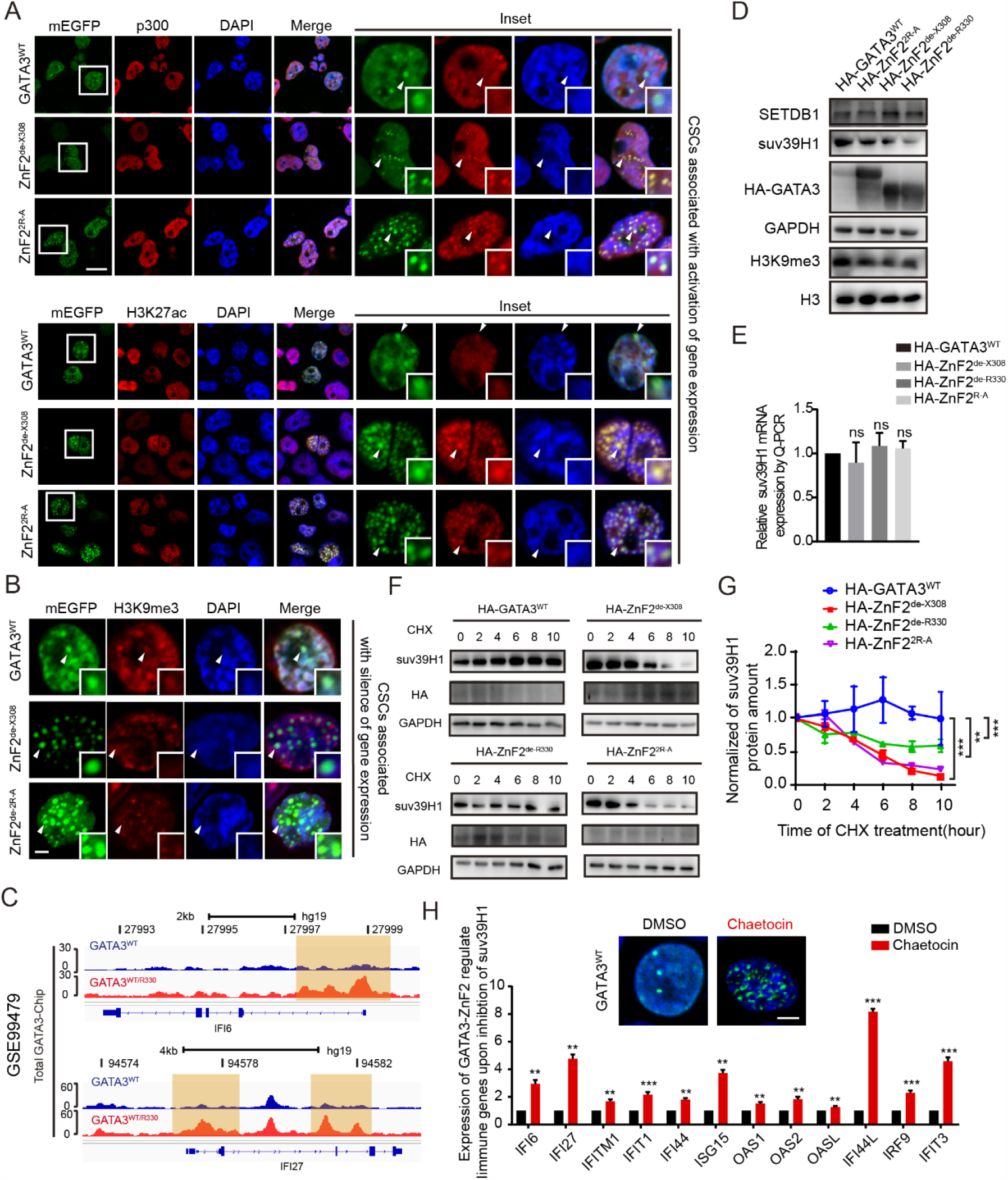
GATA3 LLPS participates in Suv39H1-regulated chromatin subcompartments modulating type I IFN signal activation. **(A)** Representative immunofluorescence confocal microscopy images by using different chromatin markers as indicated in the figures in MCF7^GATA3-KO^ with stable expression of ZnF2^2R-A^ and ZnF2^de-X308^ compared to GATA3^WT^. Green: GATA3^WT^, ZnF2^2R-A^, or ZnF2^de-X308^. Red: chromatin markers, including p300 (above), H3K27ac (below). Blue: DAPI. Scale bar,2μm. **(B)** Representative immunofluorescence confocal microscopy images by using H3K9me3 as indicated in the figures in MCF7^GATA3-KO^ with stable expression of ZnF2^2R-A^ and ZnF2^de-X308^ compared to GATA3^WT^. **(C)** Representative images of Genome Browser tracks for representative ISGs, including *IFI27* and *IFI6*, with the enriched binding of ZnF2^de-R330^ compared to wild-type GATA3 (GATA3^WT^), using publicly available chip-seq dataset with accession number: GSE99479. UCSC Genome Browser was used to show the enriched read densities of GATA3^WT^ and ZnF2-defective GATA3 mutants in or around the indicated genes in the human genome, which are colored in blue and red, respectively. Orange shading highlights the remarkable changes in the enrichment binding loci. **(D)** Representative images of western blot analysis showing the level of Suv39H1, SETDB1 and H3K9me3 in MCF7^GATA3-KO^ with stable expression of ZnF2^2R-A^, ZnF2 ^de-X308^, and ZnF2 ^de-R330^ compared to GATA3^WT^. **(E)** Suv39H1 mRNA expression levels in MCF7^GATA3-KO^ with stable expression of ZnF2^2R-A^, ZnF2 ^de-X308^, and ZnF2 ^de-R330^ compared to GATA3^WT^ by RT-qPCR. ns, not significant (t-test). **(F-G)** Suv39H1 protein levels in MCF7^GATA3-KO^ with stable expression of ZnF2^2R-A^, ZnF2 ^de-X308^, and ZnF2 ^de-R330^ compared to GATA3^WT^: (**F**) representative images of western blot analysis, (**G**) Quantification of suv39H1 protein level in the figure *D*. CHX indicates Cycloheximide treatment. GAPDH levels were used as input control. Data are mean ± s.d. **, p<0.01; ***, p<0.001 (t-test). **(H)** The impacts of Suv39H1 inhibitor-Chaetocin treatment. Above, Representative immunofluorescence confocal microscopy images showing the impact of Chaetocin treatment on GATA3 condensation on chromatin in MCF7^GATA3-KO^ with stable expression of GATA3^WT^ compared to control. Chaetocin,100 nM. Control: DMSO. Scale bar, 2 μm. Below, the impact of Chaetocin treatment on the expression of GATA3-bound ISGs as indicated in the figure in MCF7^GATA3-KO^ with stable expression of GATA3^WT^ by RT-qPCR. Data are mean ± s.d.**, p < 0.01; ***, p < 0.001 (t-test).

## Discussion

ZnF TFs play important roles in cellular and organismal functions and dysfunction in humans by controlling gene expression (Ellis et al., 2012; Jen and Wang, 2016). LLPS has emerged as a fundamental mechanism underlying the precise regulation of various biological processes, including transcriptional regulation, which is the determinant molecular event underlying both cellular and organismal phenotypic behaviors in human health and disease, including those related to cancer development (Ahn et al., 2021; Alberti, 2017; Boija et al., 2018; Bouchard et al., 2018). LLPS of ZnF TFs has been suggested to be a driving mechanism underlying their functions in controlling gene transcription by characterizing some Zn TFs, such as Oct4 and GCN4, to activate their target genes via the LLPS capacity of their IDR-containing ADs (Boija et al., 2018). GATA3 can undergo LLPS via the capacity of its IDR-containing AD (Nair et al., 2019). In addition to IDR-containing ADs, GATA3 and all other Zn TFs have ZnF-containing DBDs as their unique features. The DBD of Zn TFs has the ability to bind DNA, and ZnF plays an essential role in DBDs. However, it is largely unknown whether and how well-folded DBDs and ZnF in their DBDs can contribute to the LLPS of Zn-TFs. Here, we report that the structured DBD of GATA3 plays a master regulatory role in the LLPS of GATA3, where ZnF2 in the DBD and its R329 and R330 in ZnF2 regulate the LLPS behavior of GATA3 by providing MEIs. By demonstrating aberrant LLPS of GATA3 upon mutation of its DBD of GATA3 or the ZnF2 in its DBD or R329 and R330 in ZnF2, we claim that the DBD of a Zn TF and the ZnF in its DBD play an important role in MEI-mediated regulation of the LLPS capacity of its AD.

Cancers caused by gene mutations are a leading cause of human death, and breast cancer is the most common human cancer, with an estimated 2.3 million new cases in 2020 (Sung et al., 2021). In recent decades, extensive sequencing efforts have identified a large number of gene mutations in breast and other major cancers (Nogrady, 2020). ZnF TFs, which are frequently mutated in cancer, play critical roles in cancer initiation and progression (Jen and Wang, 2016). However, cancer-associated mutations in a given ZnF TF are myriad and lack functional and mechanistic interpretations. GATA3 is one of the three most frequently mutated genes in breast cancer (Ellis et al., 2012). Previous studies have shown that breast cancer patients with ZnF2^de-X308^ but not ZnF2^in-M293K^ have a better prognosis than those with GATA3^WT^ (Hruschka et al., 2020; Takaku et al., 2018). Another study showed that GATA3^WT^ promotes the expression of many oncogenes, such as MYB and ERα, which enhance tumor growth potential and cause a worse prognosis (Hruschka et al., 2020). Although GATA3 is considered to play a critical role in breast cancer development, its role of GATA3 in breast cancer is still under debate (Takaku et al., 2015). Here, we show that although aberrant GATA3 LLPS caused by its ZnF2 defective breast cancer-associated mutations significantly reduced the oncogenic potential of breast cancer, elevated expression of GATA3, whether mutated or not, plays an oncogenic role in breast cancer development. The ZnF-regulated LLPS of GATA3 provides a clinical phenotype-associated mechanism to unify the oncogenic role of GATA3 in the disease course of breast cancer, suggesting that LLPS of ZnF TFs regulated by their ZnFs may be useful for the functional and mechanistic interpretation of disease-associated ZnF TF mutations in many malignant settings.

The immune system is essential for human health, and its dysfunction plays a central role in the development of various human diseases including cancer (Ikeda and Togashi, 2022). GATA3 is a master regulator that controls the generation and function of immune cells, including T, B, NK, and ILC2 cells (Wan, 2014). The immune system is a highly sophisticated system that is a complex network of organs, immune cells, non-immune cells and proteins, where increasing studies have noticed essential roles of immune modulation by non-immune cells (Dainichi et al., 2021). For example, we recently reported that breast cancer cells, as non-immune cells, facilitate the formation of an immunosuppressive microenvironment via cancer-intrinsic BOMR-inhibited IRF3 (Liu et al., 2022). However, the function of GATA3 in the regulation of immunity in non-immune cells remains largely unknown. Here, we showed that aberrant GATA3 LLPS caused by its ZnF2 defective mutations promotes aberrant upregulation of immune gene expression and activation of type I IFN signaling in breast cancer cells, suggesting that ZnF2-regulated LLPS underlies the master role of GATA3 in immune regulation beyond immune cells. In cell nuclei, ZnF TFs are ubiquitously shown as microscopically visible chromatin condensates/speckles, known as CSCs, which directly localize to ZnF TFs to regulate their target gene expression (Sharma et al., 2021). CSC formation and chromatin organization are ultimately associated with ZnF TFs in shaping the transcriptional network in human health cells and disease cells, including cancer cells, owing to their oncogenic behavior (Dreijerink et al., 2017; Hruschka et al., 2020; Shin et al., 2018). The formation of CSCs is critical for the precise regulation of chromatin organization and gene transcription (Hansen et al., 2021; Narlikar, 2020). CSCs contain many proteins, including TFs, mediators, histones, and various chromatin proteins, and their formation can be modulated by various regulatory factors (Gibson et al., 2019). HP1α and H3K9me3 are two conserved hallmarks of heterochromatin (Chang et al., 2021). LLPS of HP1α is considered a driving mechanism underlying CSC formation and chromatin organization, where H3K9me3 generated by suv39H1 mediates LLPS of HP1α (Wang et al., 2019). LLPS is proposed to be one of the master mechanisms underlying ZnF TFs and CSC interactions, which are of fundamental importance for controlling the expression of genes critical for human health or disease (Jen and Wang, 2016; Karn and Emerson, 2020; Urnov, 2002). It is not surprising that Zn TFs have been implicated in LLPS-mediated CSC formation (Sharma et al., 2021). However, the underlying mechanisms of these effects are only partially understood. Here, we found that aberrant GATA3 LLPS caused by ZnF2 defective mutations remarkably altered chromatin organization, but reduced suv39H1 and H3K9me3 levels by destabilizing suv39H1. Furthermore, suv39H1 inhibition by Chaetocin increased GATA3-containing CSCs. Consistent with the remarkable increase in the expression of immune genes by GATA3 ZnF2 defective mutants compared to the wild-type protein, GATA3 ZnF2 defective mutants showed increased colocalization with transcription activation markers, including p300 and H3K27ac, supporting that the DBD of GATA3 and ZnF2 in its DBD plays an important role in controlling gene expression through an LLPS-mediated mechanism. Interestingly, Suv39H1 was recently found to be a critical factor in regulating immunity, where inhibition of Suv39H1 activates TE-encoded retroviral antigens (ERVs) to induce TE-specific cytotoxic T cell responses and activate innate immunity against tumors (Padeken et al., 2022; Pan et al., 2022). Although chronic exposure to low levels of type I IFN provides important pro-survival advantages, hyperactivated type I IFN signaling drives growth arrest and apoptosis in cancer cells (Cheon et al., 2023). Here, we found that in breast cancer cells, aberrant GATA3 LLPS caused by breast cancer-associated ZnF2 defective mutations resulted in aberrant upregulation of type I IFN signaling-related immune genes and activation of type I IFN signaling. These results suggest that aberrant GATA3 LLPS caused by breast cancer-associated ZnF2 defective mutations hyperactivate type I IFN signaling, contributing to their reduced ability to promote breast cancer growth.

In summary, our data demonstrated that ZnF2-regulated GATA3 LLPS underlies the role of GATA3 in breast cancer development and immune regulation. Since ZnFs are a common feature of ZnF TF DBDs, by describing ZnF2-regulated GATA3 LLPS as a proof-of-principle, we asserted that the LLPS of ZnF TFs regulated by ZnF in their DBDs is an important mechanism underlying the multifaceted function of ZnF TFs in human health and diseases such as cancer.

## Materials and Methods Cell lines and cell culture

The cells were maintained in accordance with the guidelines of the American Type Culture Collection. Human HEK293T and HeLa cells were cultured in DMEM supplemented with 10% (v/v) FBS, 100 U/mL penicillin, and 100 mg/mL streptomycin. MCF7, T47D, and 4T1 cells were cultured in RPMI1640 supplemented with 10% (v/v) FBS, 100 units/mL penicillin, and 100 mg/mL streptomycin. All the cells were cultured at 37 °C in a humidified atmosphere of 95% air and 5% CO_2_. All cells were verified by periodic morphological checks and mycoplasma detection.

### Mouse model and experiments details

Mice were housed in AAALAC-accredited facilities and maintained on a 12□h light:12□h dark-light cycle at room temperature (21–23□°C) and 40–60% humidity. The mice were housed separately by sex (n□=□5 per cage) in corncob bedding with filter-top lids. The maximal tumor measurement permitted by the IACUC was 20□mm in diameter. Tumor volumes and other endpoints did not exceed the limits permitted by the IACUC. All mice were maintained under specific pathogen-free conditions and were housed in individually ventilated cages.

4T1 cells transduced with GATA3 constructs were resuspended in 100□ µL PBS and injected into the tail vein of 7-week-old BALB/C mice. Tumor size was measured in the two longest dimensions using a Vernier caliper. The tumor volume (V) was calculated using the formula V□=□D_1_(D_2_)^2^/2, where D_1_ and D_2_ are the long and short dimensions, respectively. Twenty days after transplantation, the mice were humanely euthanized to collect tumors.

## METHOD DETAILS

### RNA extraction and qRT-PCR

Total RNA was extracted using the XGEN Reagent. All experiments were performed according to the manufacturer’s instructions. Briefly, RNAs was reverse-transcribed into cDNA using HiScript II Q-RT Super Mix (Vazyme), and qPCR was performed using ChamQ SYBR Q-PCR Master Mix (Vazyme). Relative mRNAs and mRNAs expression levels were calculated using the 2^−ΔΔCt^ method. GAPDH served as an internal control. Primer sequences for RT–qPCR are listed in *SI Appendix* (Table S1).

### Vector construction

pmEGFP-N2 was mutated to pEGFP-N2 (monomeric EGFP, with the A206K mutation of EGFP). The GATA3-WT vector was purchased (Youbia). We initially designed primers to generate different regions of GATA3-WT (443 aa), AD (1-260aa), DBD (261-360aa), C (361-443aa), △AD (261-443aa), △DBD (1-260aa+361-443aa) and △C (1-360aa), then these truncations subclone into the pmEGFP-N2. GATA3 and truncations were generated by PCR and subcloned into the pET-MBP-3C vector. Mutant GATA3-mEGFP containing various truncations and mutations, such as ZnF2^in-G360^ and ZnF2^de-R330^ were generated using PCR and similarly cloned into the above vectors. The ZnF2^de-X308^ splice mutant is a genome splice protein generated by GATA3-WT, excluding the TCTGCAG sequence from S308.

Other GATA3 point mutations were also generated by GATA3-WT, such as ZnF2^in-M293K^, ZnF1^2R-A^ (R275R276-A275A276), ZnF2^2R-A^(R329R330-A329A330) and GATA3^4R-A^ (R275R276R329R330-A275A276A329A330). The MBP-GATA3 plasmids were PCR using pmEGFP-N2-GATA3 variants to be inserted at the BamHI and NotI sites in the pET-MBP-3C construct. HA-tag GATA3 mutants and WT construct subclones to pCDH-CMV-MCS-EF1-copGFP-T2A-Puro and BamHI and NheI sites were used in pCDH-CMV-MCS-EF1-copGFP-T2A-Puro. The primers used are listed in the SI Appendix, Table S2.

### Recombinant protein expression and purification

For recombinant protein expression in E. coli, full-length GATA3, and truncations were cloned into the pET-MBP-3C vector with an N-terminal His-MBP tag. For MBP-GATA3, WT/mutants-mEGFP were clones from pmEGFP-GATA3 WT/ mutants to the pET-MBP-3C vector with a C-terminal mEGFP tag and HRV 3C protease cleavable N-terminal His-MBP tag.

GATA3 and its mutants were expressed at 18 °C for 16 h with 0.2 mM IPTG induction. All the proteins were expressed in Escherichia coli BL21 (DE3) cells. Cells were grown to OD 0.6-0.8 at 37 °C in LB media with 100 mg/ml ampicillin. E. coli cells expressing protein were induced with 0.1-0.3mM IPTG at 16 °C for 18 h, followed by sonication in lysis buffer (50 mM Tris-HCl pH 7.5, 500 mM NaCl, 1 mM PMSF). The cells were maintained at 80% of 300 W power, on 5 s, off 10 s, and the remaining buffer was clarified. These proteins were first purified using Ni-affinity chromatography, and then further purified using Superdex 200 Increase 10/300 GL. These proteins were flash-frozen in liquid nitrogen and stored in 50 mM Tris-HCl pH 7.5, 500 mM NaCl, 1 mM DTT, and 10% glycerol at -80 °C. Purified proteins were examined by SDS–PAGE followed by Coomassie blue staining. Protein concentration was determined by Nanodrop measurement for OD 562 and calculated using the extinction coefficient provided by ExPASy ProtParam (https://web.expasy.org/protparam/) and validated by comparison with Coomassie Blue staining of known concentrations of BSA.

### Generation of MCF7^GATA3-KO^ and stable HA-GATA3 cell lines

Briefly, guide RNA was designed using the Dr. Feng Zhang Laboratory web tool (crispr.mit.edu). Double-stranded oligonucleotides were inserted into lentiCRISPR-v2(Addgene Plasmid 52961). HEK293T cells were co-transfected with 2 μg lentiCRISPR-v2 vector and lentivirus packaging plasmid1.5 μg pSPAX2 and 0.5 μg pMD2. G, plasmid using Lipofectamine 2000 reagent according to the manufacturer’s instructions. After incubation for 48h, the collection virus-infected MCF7 cells, after 48h incubation, 2ug/ml puromycin was used to select positive cells. The positive cell limit dilution was then spread into 96-well plates for culture. After 2 weeks, we observed a significant colon in 96-well plates, and these cells were chosen to detect GATA3 by western blotting. Stable HA-GATA3 cells expressing GATA3 variants were generated by retroviral infection of MCF7^GATA3-KO^ and 4T1 cells using pCDH-CMV-MCS-EF1-copGFP-T2A-Puro. Briefly, HA-tag labeled GATA3 mutants and WT construct were subcloned into the pCDH-CMV vector, and then transfected into HEK293T cells for 48h. Collection virus to infection cells, after 48h incubation, 2ug/ml puromycin was used to choose stable expression GATA3 WT and mutant cells 14 days. The primers used are listed in the SI Appendix, Table S3.

### In vitro droplets formation

In vitro LLPS experiments were performed at room temperature unless otherwise indicated. For the droplet formation assay, the purified GATA3 WT or mutant proteins without the mEGFP-tag were mixed with a buffer containing 50 mM Tris pH 7.4 and incubated for 2 h at room temperature after adding proteinase 3C at different NaCl concentrations. The samples were observed under a DIC microscope using a Leica microscope with a 40x objective. PreScission protease 3C cleaves N-terminal His-MBP-tag for recombinant protein. MBP-mEGFP and MBP-GATA3 variants of mEGFP were also performed at room temperature with a buffer containing 50 mM Tris pH 7.4 and incubated for 2 h at room temperature after adding proteinase 3C at different NaCl concentrations. Finally, 10μL of each sample was pipetted into a glass dish and imaged using a Nico TIAR microscope. Under the same conditions, 10 μL of each sample was added to a 384-well white polystyrene plate with a clear, flat bottom. For protein and DNA mixing, GATA-DNA or non-GATA-DNA was incubated before GATA3 proteins were cleaved by PreScission protease 3C, and then 10 μL of each sample was added to a 384-well white polystyrene plate with a clear flat bottom and images. The DNA sequences are listed in the SI Appendix, Table S4.

### Turbidity assays

The turbidity of the phase-separated samples was measured based on the optical absorption at 600 nm using a TECAN Infinite F PLEX Pro microplate reader. Turbidity measurements of GATA3 proteins under 3C cleavage were recorded on a spectral scanning multimode reader (Thermo Fisher) using flat-bottom and low-volume 384-well plates (Corning).

### Western blot

Cells were lysed in RIPA lysis buffer (Beyotime, P10030B) supplemented with 1 mM 1 mM phenylmethylsulfonyl fluoride (PMSF). Protein concentrations were quantified using the BSA protein assay kit. Briefly, 30 μg of the protein extract was loaded and separated using SDS-PAGE gels. Blotting was performed according to the standard protocols. All 0.45μm and 0.22 μm PVDF membranes (Millipore) were used. Membranes were blocked for 1 h in blocking buffer (5% BSA in TBST), followed by incubation with primary antibody probes at 4°C. Membranes were incubated with HRP-conjugated secondary antibodies after three washes with PBST. The signals were detected using BeyoECL Plus (Beyotime, P0018S). All antibodies used for immunoblotting are listed in *SI Appendix*, Table S5.

### Immunofluorescence staining and live-cell imaging

Immunofluorescence analysis was performed according to standard protocols. Briefly, cells were washed with PBS, fixed with ice-cold methanol for 15□min, and then cultured in Triton-100 for 30 min. Endogenous p300, H3K27ac, and H3K9me3 bound to Alexa 594 were also detected with mEGFP-tag GATA3 in MCF7^GATA3-KO^ cells. The original image resolution was 1024 x 1024 pixel^2^. Cells were incubated with rabbit antibodies p300(Abcam, ab275378,1:200), H3K27ac (Abcam, ab4729,1:200) and H3K9me3(Abcam, ab8898:1:200) at 4 °C overnight, followed by incubation with Alexa Fluor 594 goat anti-rabbit antibody (Abcam, ab150080,1:200) for 2 h at room temperature. Slides were further counterstained with DAPI.

For live-cell imaging, cells were cultured in 35□mm glass-bottom dishes (Corning) and used for imaging on a Zeiss LSM710 confocal microscope supported by a Chamlide TC temperature, humidity, and CO_2_ chamber. High content image. HEK293T cells expressing GATA3-WT, GATA3 truncations (N, C, DBD, △N, △C, and △DBD), and GATA3 mutants (ZnF2^in-2R-A^, ZnF2^de-2R-A^, and GATA3^4R-A^) were seeded in 24-well glass bottom plates (Sofa P24-1.5H-N), and images were obtained using the NIS-element cell imaging analysis system (Nico TIAR). The 100x objective lens was used for each condition. Data were analyzed using images collected from 10 representative fields in each group. Single cells were identified using DAPI as a reference, while spot quantification was performed based on the area and intensity of the spots through the mEGFP channel. MCF7GATA3-KO or T47D cells expressing GATA3WT and GATA3 mutants (ZnF2in mutants and ZnF2de mutants) and images were obtained using the NIS-element cell imaging analysis system (Nico TIAR). The resolution of the original image was 1024 x 1024 pixel^2^.

### Fluorescence Recovery After Photobleaching (FRAP)

The FRAP assay was conducted using the FRAP module of a Zeiss LSM710/810 ZEN blue confocal microscopy system. GATA3-mEGFP proteins were bleached using a 488 nm laser beam. Bleaching was focused on a circular region of interest (1-2μm) using a 100% laser power, and after approximately 5 s, time-lapse (2 s per image) images were collected. For GATA3 droplet FRAP in live cells, Leica STELLARIS STED or Zeiss LSM710 was used for the assay. The Zeiss LSM710 bleaching set consisted of in vitro and in vivo proteins. For Stellaris STED, the bleaching time was 3 s and the time interval was 2 s per image. Time images remain 10 min. The fluorescence intensity was measured using a ZEN Blue or Leica system. The background intensity was subtracted, and values were reported relative to the pre-bleaching time points. The percentage recovery for each time point was calculated as (I − I_min_)/ (I_0_ − I_min)_ × 100%, where I, I_min_, and I_0_ are the normalized intensity for each time point, minimum (0 s after bleaching), and initial (before bleaching) intensities, respectively. FRAP recovery curves were fitted to generate the half-time recovery for each bleached condensate, which was used to calculate the mean and s.d. of the half-time. GraphPad Prism was used to plot and analyze the FRAP results.

### Cell growth assay

For cell growth, GATA3 variants stably expressing MCF7 cells were seeded in a 12-well plate at a density of 1 × 10^5^ cells/well and the number of cells number (1,2,3,4 day). GraphPad Prism was used to plot and analyze the results.

### EMSA assay

DNA binding of purified GATA3^WT^ and mutants was assessed by incubating increasing concentrations of GATA3 with 1μM biotinylated DNA for 30 min at RT, in HEPES 50 mM, pH 7.5, 150 mM NaCl, 1 mM DTT, and 10 mM MgCl_2_. Complexes were then separated on TBE 4%–12% gels at 100V in TBE buffer (0.5 ×) and transferred onto a nylon membrane.

### RNA library constructing and sequencing

Total RNA (1 μg total RNA was used for library preparation. Poly (A) mRNA was isolated using oligo (dT) beads. mRNA fragmentation was performed using divalent cations and high temperatures. Priming was performed using Random Primers. First-strand and second-strand cDNA were synthesized. The purified double-stranded cDNA was then treated to repair both ends and a dA-tailing was added in one reaction, followed by T-A ligation to add adaptors to both ends. Size selection of Adaptor-ligated DNA was then performed using clean DNA beads. Each sample was then amplified by PCR using P5 and P7 primers, and the PCR products were validated. Libraries with different indices were multiplexed and loaded on an Illumina HiSeq/Illumina Novaseq/MGI2000 instrument for sequencing using a 2 × 150 paired-end (PE) configuration according to the manufacturer’s instructions.

Differential expression analysis was performed using the DESeq2 Bioconductor package, a model based on negative binomial distribution. estimates of dispersion and logarithmic fold changes incorporate data-driven prior distributions. For volcano plots, log2FC>1 or<-1, p<0.05, were used to screen for differential gene expression, and the Sanger box tool (http://sangerbox.com/) was used to draw volcano plots. For GSEA, WebGestalt (WEB-based GEne Set Analysis Toolkit) was used.

## QUANTIFICATION AND STATISTICAL ANALYSIS

All data are expressed as mean ± s.d. and analyzed using GraphPad Prism 7.0 software. For experiments consisting of only two groups, the data were evaluated using Student’s t-test (two tails). For experiments involving multiple groups, the data were analyzed using one-way analysis of variance. The log-rank test was used for survival analysis. Statistical significance was set at p < 0.05. GraphPad Prism 7 was used to employ unpaired or paired t-tests or log-rank tests where appropriate. p-values are denoted as *, p < 0.05; **, p < 0.01; ***, p < 0.001; NS, not significant.

### Data Availability

Additional procedures are described in the SI Appendix (Supplementary Information). All study data are included in the article and the SI Appendix.

## Acknowledgments

This work was supported by the National Natural Science Foundation of China (81974447), the Natural Science Foundation of Jiangsu Province (SBK2020010058), and the Priority Academic Program Development of Jiangsu Higher Education Institutions.

## Notes

### Competing Interest Statement

The authors have declared no competing interest.

### Summary of Updates

result 1-7

