## Supplementary material for "GATA3 ZnF2-defective mutant condensation underlies type I IFN-activating in breast cancer": s1-s7

**This PDF file includes:**

Figures S1 to S7
Tables S1-S5

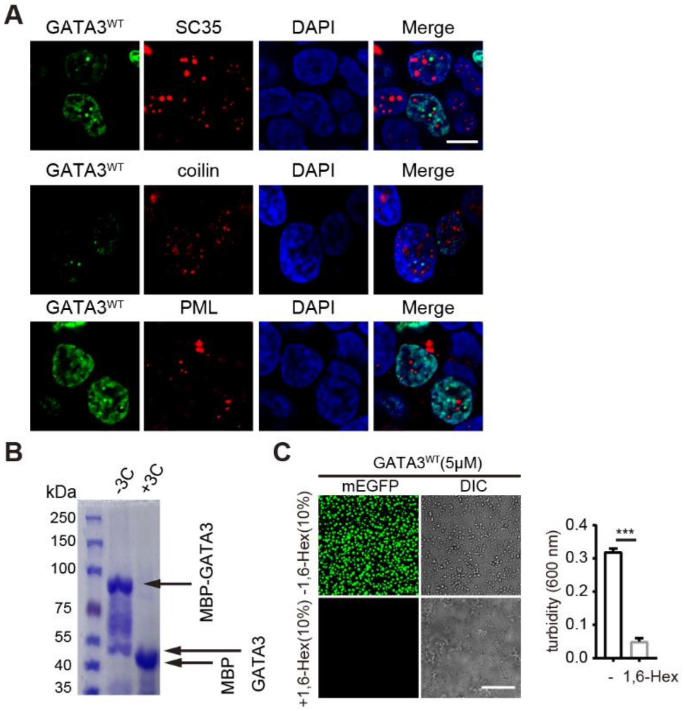


**Fig. S1. Nuclear GATA3 condensations and the impact of 1,6-Hex on the LLPS of purified GATA3 protein *in vitro*. (A)** Representative confocal microscopy images of HEK293T cells with stable expression of the GATA3$^{WT}$-mEGFP gene. PML, coilin or SC35 labeled by immunofluorescence were counterstained with DAPI as indicated. Scale bar, 5 μm. **(B)** Coomassie blue staining of 10μM purified recombinant MBP-GATA3 proteins with and without 3C precession protease treatment. **(C)** In vitro LLPS assays of MBP-GATA3-mEGFP after 3C precession protease treatment. Left, representative images of droplets formed by 5 μM MBP-GATA3-mEGFP with and without 10% 1,6-Hex treatment. Right, quantification of solution turbidity of 5 μM MBP-GATA3-mEGFP with and without 10% 1,6-Hex treatment by measuring optical density at 600 nm (OD600; means ± s.d., n = 3 experiments). Data are mean ± s.d. ***, p<0.001 (t-test).

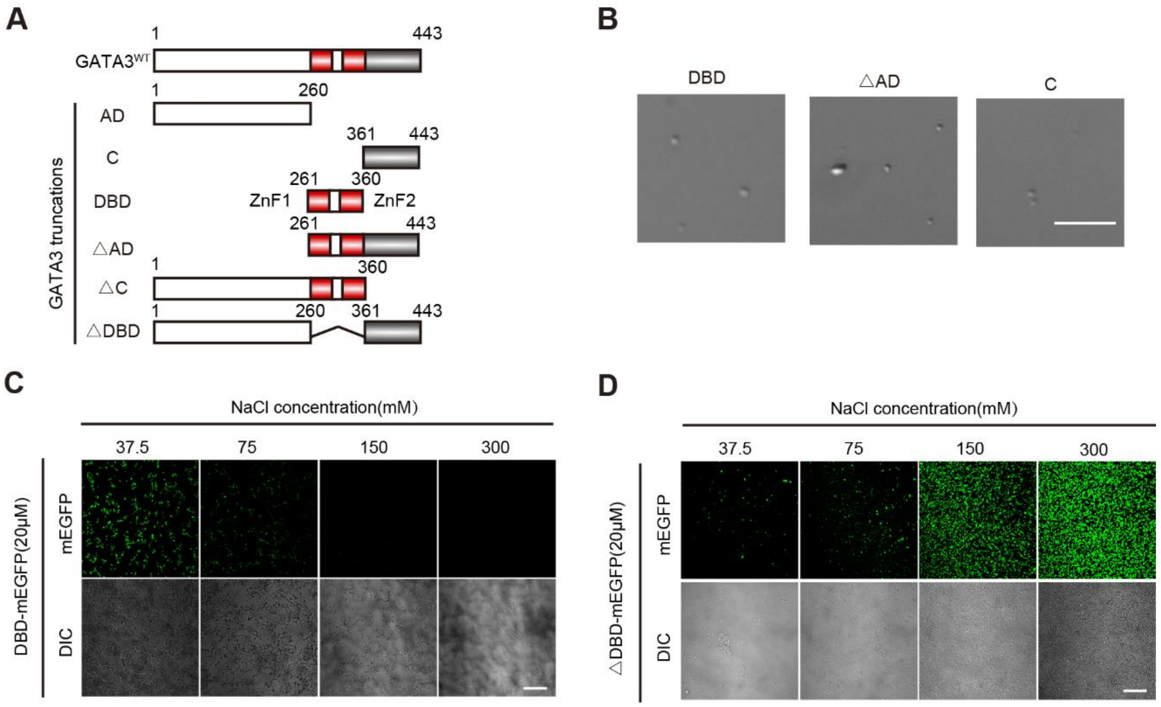


**Fig. S2. GATA3 truncations and the effect of salt concentration on the LLPS of purified DBD and △DBD proteins *in vitro*. (A)** Diagrams for the domain structure of GATA3 truncations. **(B)** Representative images of droplets formed by 10 μM purified different MBP-GATA3 domain truncated proteins without mEGFP tag after cleavage of MBP by 3C protease treatment. Scale bar, 20 μm. **(C-D)** Representative images of the LLPS of 20μM purified mEGFP-tagged DBD (**C**) and mEGFP-tagged △DBD (**D**) proteins with increasing of NaCl concentration from 37.5 mM to 300 μM. Scale bar, 10 μm.

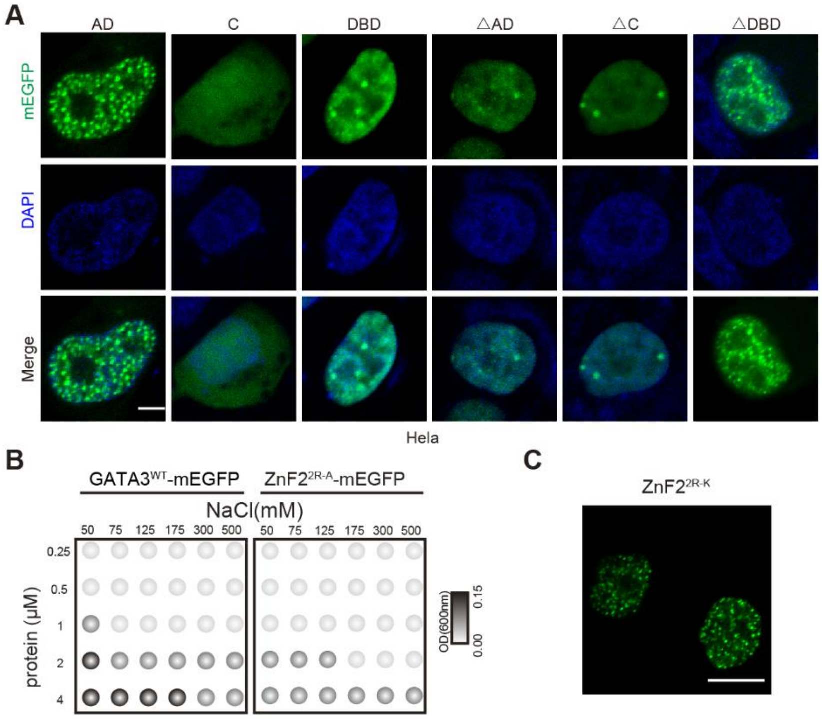


**Fig. S3. LLPS ability of purified ZnF2$^{2R-A}$ protein compared to GATA3$^{WT}$ *in vitro.* (A)** Representative confocal microscopy images of Hela cells with ectopic expression mEGFP-tagged truncations. Scale bar, 2 μm. **(B)** Phase plots summarizing the LLPS of purified GATA3$^{WT}$-mEGFP and ZnF2$^{2R-A}$-mEGFP proteins with the concentrations ranging from 0.25–4 μM with increasing NaCl concentration from 50 to 500 mM. White dots, no phase separation; gray and black dots, phase separation. The LLPS ability of GATA3$^{WT}$-mEGFP and ZnF2$^{2R-A}$-mEGFP under different conditions was Gray-coded on the basis of droplet turbidity measured at OD600 after the proteins were incubated with phase separation buffer at room temperature for 120 min. **(C)** Representative confocal microscopy images of HEK293T cells with ectopic expression of mEGFP-tagged GATA3 mutants ZnF2$^{2R-K}$ as indicated in the figure. Scale bar, 5 μm.

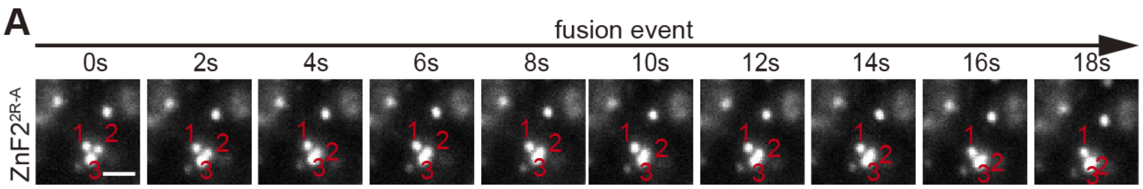


**Fig. S4. ZnF2$^{2R\text{-}A}$ condense fusion event in cells. (A)** Representative images showing the fusion event of ZnF2$^{2R\text{-}A}$-mEGFP in HEK293T cell. Scale bar, 1 μm. Three droplets are indicated by red numbers, 1, 2, and 3, where droplets 2 and 3 were fused over the indicated time.

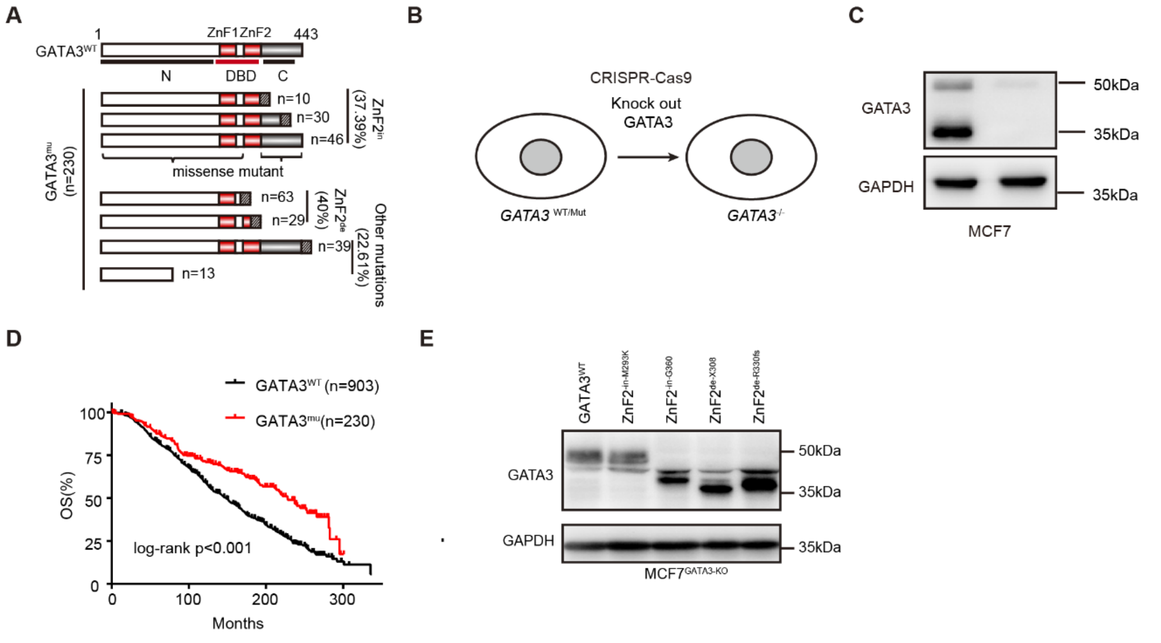


**Fig. S5. Analysis of breast cancer-associated GATA3 mutations and generation of $MCF7^{GATA3\text{-}KO}$ cells from MCF7 by knocking out GATA3 using CRISPR-Cas9 to further generate breast cancer cells with stable overexpression of $GATA3^{WT}$ or GATA3 mutants. (A)** Summary and classification of GATA3 mutations identified in clinical breast cancer samples. Among all of GATA3 mutants, 40% are ZnF2 defective mutants ($ZnF2^{de}$), 37.39% are ZnF2 intact mutants ($ZnF2^{in}$), and 22.61% are other mutants. **(B-C)** Generation of $MCF7^{GATA3\text{-}KO}$ cells from MCF7 by knocking out GATA3 using CRISPR-Cas9: (**B**) a diagram explaining how to knock out GATA3 in MCF7 cells, (**C**) Representative images of western blot analysis confirming that GATA3 are successfully knocked out in $MCF7^{GATA3\text{-}KO}$ cells using parental MCF7 cells as controls. **(D)** Overall survival analysis of breast cancer patients with GATA3 mutations ($GATA3^{mu}$) compared to those with wild type GATA3 ($GATA3^{WT}$). n, the number of patients. **(E)** Representative images of western blot analysis confirming $MCF7^{GATA3\text{-}KO}$ cells with stable overexpression of $GATA3^{WT}$ or GATA3 mutants as indicated. GAPDH levels were used as an input control.

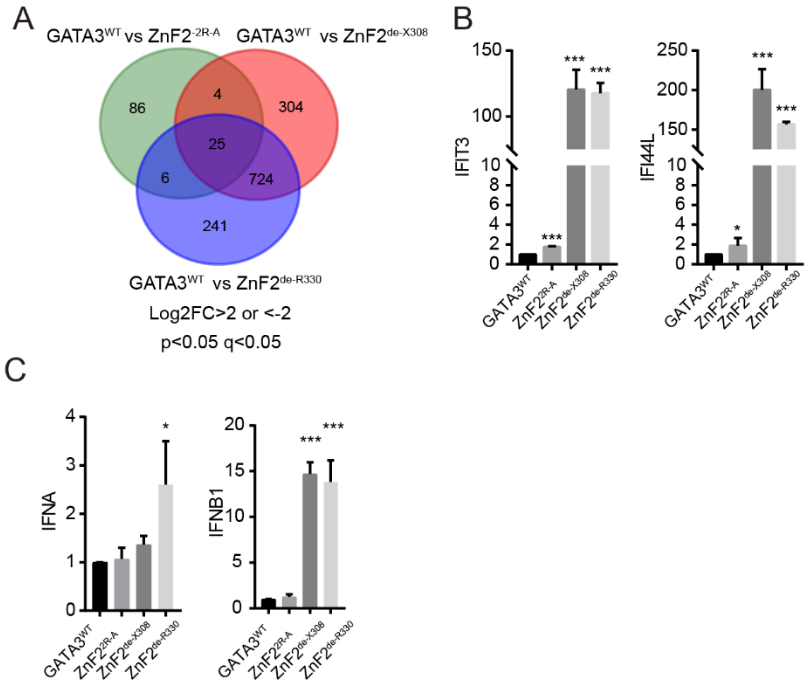


**Fig. S6. ZnF2-defective GATA3 mutants increase the expression of ISGs compared to wild-type GATA3. (A)** Venn diagram summarizing and comparing the differentially expressed genes between ZnF2$^{2R\text{-}A}$, ZnF2$^{de\text{-}X308}$, and ZnF2$^{de\text{-}X308}$ compared to GATA3$^{WT}$. **(B)** The expression of indicated ISGs (*IFIT3* and *IFI44L*) in MCF7$^{GATA3\text{-}KO}$ with stable expression of ZnF2$^{2R\text{-}A}$, ZnF2 $^{de\text{-}X308}$, and ZnF2 $^{de\text{-}R330}$ compared to GATA3$^{WT}$ by RT-qPCR in supplemented to Fig. 6E. Data are mean ± s.d. *, $p < 0.05$; **, $p < 0.01$; ***, $p < 0.001$ (t-test). **(C)** The expression of (*IFNA* and *IFNB*) in MCF7$^{GATA3\text{-}KO}$ with stable expression of ZnF2$^{2R\text{-}A}$, ZnF2 $^{de\text{-}X308}$, and ZnF2 $^{de\text{-}R330}$ compared to GATA3$^{WT}$ by RT-qPCR.

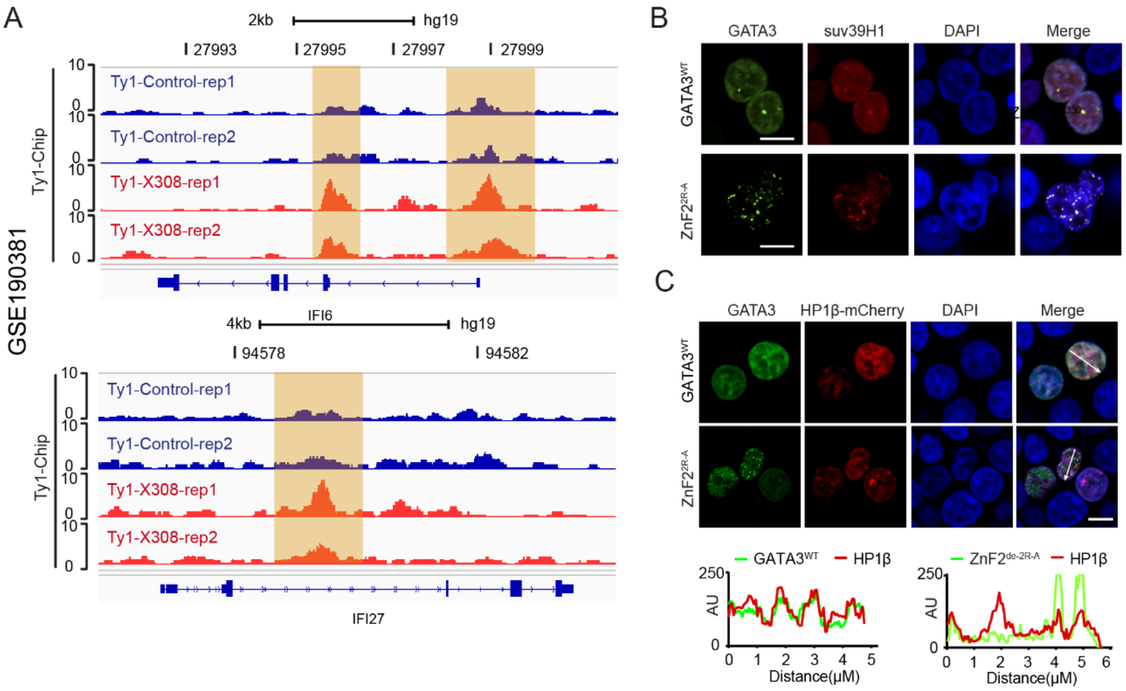


**Fig. S7. ZnF2-defective GATA3 LLPS mutants increase the expression of ISGs regulated by Suv39H1. (A)** ZnF2$^{de\text{-}X308}$ compared to GATA3$^{WT}$ using publicly available chip-seq dataset with accession number: GSE190381. UCSC Genome Browser was used to show the enriched read densities of GATA3$^{WT}$ and ZnF2-defective GATA3 mutants in or around the indicated genes in the human genome, which are colored in blue and red, respectively. Orange shading highlights the remarkable changes in the enrichment binding loci. **(B)** Representative confocal microscopy images of HEK293T cells with ectopic expression mEGFP-tagged GATA3$^{WT}$ and ZnF2$^{2R\text{-}A}$ with Suv39h1-mCherry. Scale bar, 5 μm. **(C)** Above, representative confocal microscopy images of HEK293T cells with ectopic expression mEGFP-tagged GATA3$^{WT}$ and ZnF2$^{2R\text{-}A}$ with HP1β-mCherry. Scale bar, 5 μm. Below,

**Table S1**

| Plasmid name | Forward primer sequence | Reverse primer sequence |
|---|---|---|
| p-mEGFP-N2-GATA3$^{WT}$ | CTAGCTAGCATGGAGGTGAC GGCGGACCAGCCG | CCGGAATTCACCCATGGCGG TGACCATGCT |
| p-mEGFP-N2-GATA3-AD | CTAGCTAGCATGGAGGTGAC GGCGGACCAGCCG | CCGGAATTCCAGCAGGCTGC TGGGCGGGAAGAG |
| p-mEGFP-N2-GATA3-DBD | CTAGCTAGCATGGGCGGCTC CCCCACCGGCTTCGGA | CCGGAATTCGCCTTCCTTCTT CATAGTCAGGGG |
| p-mEGFP-N2-GATA3-C | CTAGCTAGCATGATCCAGAC CAGAAACCGAAAAATG | CCGGAATTCACCCATGGCGG TGACCATGCT |
| p-mEGFP-N2-GATA3-△AD | CTAGCTAGCATGGGCGGCTC CCCCACCGGCTTCGGA | CCGGAATTCACCCATGGCGG TGACCATGCT |
| p-mEGFP-N2-GATA3-△C | CTAGCTAGCATGGAGGTGAC GGCGGACCAGCCG | CCGGAATTCGCCTTCCTTCTT CATAGTCAGGGG |
| p-mEGFP-N2-GATA3-△DBD | CTAGCTAGCATGGAGGTGAC GGCGGACCAGCCG | CCGGAATTCACCCATGGCGG TGACCATGCT |
| p-mEGFP-N2-ZnF2$^{in-M293K}$ | CCTGCGGGCTCTATCACAAA AAGAACGGACAGAACCGGCC CC | GGGGCCGGTTCTGTCCGTTC TTTTTGTGATAGAGCCCGCA |
| p-mEGFP-N2-ZnF2$^{in-G360}$ | CTAGCTAGCATGGAGGTGAC GGCGGACCAGCCG | CCGGAATTCGCCTTCCTTCTT CATAGTCAGGGG |
| p-mEGFP-N2-ZnF2$^{de-X308}$ | CTAGCTAGCATGGAGGTGAC GGCGGACCAGCCG | CCGGAATTCGGGGTCTGTTA ATATTGTGAA |
| p-mEGFP-N2-ZnF2$^{de-R330}$ | CTAGCTAGCATGGAGGTGAC GGCGGACCAGCCG | CCGGAATTCCCTCCTCCAGA GTGTGGTTGTGGT |
| p-mEGFP-N2-ZnF1$^{2R-A}$ | CGACCCCACTGTGGGCCGCA GATGGCACGGGA | TCCCGTGCCATCTGCGGCCC ACAGTGGGGTCG |
| p-mEGFP-N2-ZnF2$^{2R-A}$ | CACCACCACCACCCTCTGGG CAGCAAACGCTAATGGGGAC CC | GGGTCCCCATTAGCGTTTGC TGCCCAGAGGGTGGTGGTG |
| PCDH-HA-GATA3$^{WT}$ | CTAGCTAGCATGTACCCATAC GACGTCCCAGACTACGCTAT GGAGGTGACGGCGGACCAG CCG | ATTTGCGGCCGCTCAACCCA TGGCGGTGACCATGCT |

| | | |
|---|---|---|
| PCDH-HA-ZnF2$^{\text{in-G360}}$ | CTAGCTAGCATGTACCCATACGACGTCCCAGACTACGCTATGGAGGTGACGGCGGACCAGCCG | ATTTGCGGCCGCTCAGCCTTCCTTCTTCATAGTCAGGGG |
| PCDH-HA-ZnF2$^{\text{de-X308}}$ | CTAGCTAGCATGTACCCATACGACGTCCCAGACTACGCTATGGAGGTGACGGCGGACCAGCCG | ATTTGCGGCCGCTCAGGGGTCTGTTAATATTGTGAA |
| PCDH-HA-ZnF2$^{\text{de-R330}}$ | CTAGCTAGCATGTACCCATACGACGTCCCAGACTACGCTATGGAGGTGACGGCGGACCAGCCG | ATTTGCGGCCGCCTACCTCCTCCAGAGTGTGGTTGTGG |
| PET-MBP-GATA3$^{\text{WT}}$ | CTGTACTTCCAATCCAATATTATGGAGGTGACGGCGGACCAGC | TGGTGGTGGTGGTGGTGCTCGAGTTAACCCATGGCGGTGACCATGCT |
| PET-MBP-GATA3-AD | CTGTACTTCCAATCCAATATTATGGAGGTGACGGCGGACCAGC | TGGTGGTGGTGGTGGTGCTCGAGTTACAGCAGGCTGCTGGGCG |
| PET-MBP-GATA3-DBD | CTGTACTTCCAATCCAATATTATGGGCGGCTCCCCCACCGGCT | TGGTGGTGGTGGTGGTGCTCGAGTTAGCCTTCCTTCTTCATAGT |
| PET-MBP-GATA3-C | CTGTACTTCCAATCCAATATTATGATCCAGACCAGAAACCGAA | TGGTGGTGGTGGTGGTGCTCGAGTTAACCCATGGCGGTGACCATGCT |
| PET-MBP-GATA3-△AD | CTGTACTTCCAATCCAATATTATGGGCGGCTCCCCCACCGGCT | TGGTGGTGGTGGTGGTGCTCGAGTTAACCCATGGCGGTGACCATGCT |
| PET-MBP-GATA3-△ C | CTGTACTTCCAATCCAATATTATGGAGGTGACGGCGGACCAGC | TGGTGGTGGTGGTGGTGCTCGAGTTAGCCTTCCTTCTTCATAGT |
| PET-MBP-GATA3-△ DBD | CTGTACTTCCAATCCAATATTATGGAGGTGACGGCGGACCAGC | TGGTGGTGGTGGTGGTGCTCGAGTTAACCCATGGCGGTGACCATGCT |
| PET-MBP-GATA3$^{\text{WT}}$-mEGFP | CTGTACTTCCAATCCAATATTATGGAGGTGACGGCGGACCAGC | TGGTGGTGGTGGTGGTGCTCGAGTTACTTGTACAGCTCGTCCAT |
| ZnF1$^{\text{2R-A}}$ | CGACCCCACTGTGGGCCGCAGATGGCACGGGA | TCCCGTGCCATCTGCGGCCCACAGTGGGGTCG |
| ZnF2$^{\text{2R-A}}$ | ACCACAACCACACTCTGGGCAGCAAATGCCAATGGGGA | TCCCCATTGGCATTTGCTGCCCAGAGTGTGGTTGTGGT |

| | | |
|---|---|---|
| M293K | CCTGCGGGCTCTATCACAAAA AGAACGGACAGAACCGGCCCC | GGGGCCGGTTCTGTCCGTT CTTTTTGTGATAGAGCCCG CAGG |

**Table S2. Primers used in this paper for Cas-9.**

| Cas-9 | Forward primer sequence | Reverse primer sequence |
|---|---|---|
| GATA3-gRNA1 | CACCGGACGGCGGACCAGCCGCGCT | AAACAGCGCGGCTGGTCCGCCGTCC |
| GATA3-gRNA1 | CACCGCACCACCCCGCCGTGCTCAA | AAACTTGAGCACGGCGGGGTGGTGC |

**Table S3. Primers used in this paper for DNA and GATA3 protein.**

| DBD binding | Forward primer sequence | Reverse primer sequence |
|---|---|---|
| no-GATA-DNA | AATGTCAAACTTTTAAGACG | CGTCTTAAAAGTTTGACATT |
| GATA-DNA | AATGTCCATCTGATAAGACG | CGTCTTATCAGATGGACATT |

**Table S4. Primers used in this paper for Q-PCR.**

| Gene name | Forward primer sequence | Reverse primer sequence |
|---|---|---|
| GAPDH | AACATCATCCCTGCCTCTACTGG | GTTTTTCTAGACGGCAGGTCAGG |
| β-actin | AGAGCTACGAGCTGCCTGAC | AGCACTGTGTTGGCGTACAG |
| OAS1 | TGTCCAAGGTGGTAAAGGGTG | CCGGCGATTTAACTGATCCTG |
| OAS2 | AGGTGGCTCCTATGGACGG | TTTATCGAGGATGTCACGTTGG |
| OASL | CTGATGCAGGAACTGTATAGCAC | CACAGCGTCTAGCACCTCTT |
| IFIT1 | TTGATGACGATGAAATGCCTGA | CAGGTCACCAGACTCCTCAC |
| IFITM1 | GGTCCCTGTTCAACACCCTC | GTCACAGAGCCGAATACCAGT |
| IFIT3 | GGGCAGACTCTCAGATGCTC | TCAAAACACACCTTCGCCCT |
| IFI44 | ATGGCAGTGACAACTCGTTTG | TCCTGGTAACTCTCTTCTGCATA |
| IFI44L | TGGTTGAAAGATGCAGCCGT | ACCATAGGGAATCATTTGGCTCT |
| IFI6 | GGTCTGCGATCCTGAATGGG | TCACTATCGAGATACTTGTGGGT |
| IFI27 | TGCTCTCACCTCATCAGCAGT | CACAACTCCTCCAATCACAACT |
| IRF9 | AGCTTGAGAGGGGCATCCTA | TATGCAACACACAAGCGCAG |
| ISG15 | CGCAGATCACCCAGAAGATCG | TTCGTCGCATTTGTCCACCA |
| suv39H1 | GCATCGCTTTCTTTGCCACA | AGGTATTTGCGGCAGGACTC |

**Table S5. Antibody used in this paper for WB or IF.**

| Antibody | Cat# | used |
|---|---|---|
| GFP | ABclonal:AE012 | WB(1:1000) |
| GAPDH | ABclonal:AC033 | WB(1:1000) |
| GATA3 | abcam: ab199428 | WB(1:2000) |
| IRF3 | abcam: ab68481 | WB(1:2000) |
| p-IRF3-S386 | CST: 37829 | WB(1:2000) |
| p-IRF3-S396 | Affinity: AF2436 | WB(1:2000) |
| TBK1 | ABclonal:A3458 | WB(1:1000) |
| p-TBK1-S172 | ABclonal:AP1418 | WB(1:1000) |
| RIG-I | ABclonal: A22478 | WB(1:1000) |
| suv39H1/KMT1A | abcam: ab12405 | WB(1:2000) |
| SETDB1 | abcam:ab107225 | WB(1:1000) |
| H3 | abcam:ab1791 | WB(1:2000) |
| H3K9me3 | abcam:ab8898 | WB(1:2000) IF(1:200) |
| H3K27ac | abcam:ab4729 | IF(1:200) |
| HA | abcam:ab9110 | WB(1:2000) |
| p300 | abcam:ab275378 | IF(1:200) |
